# Molecular characterization of Rft1, an ER membrane protein associated with congenital disorder of glycosylation RFT1-CDG

**DOI:** 10.1101/2024.04.03.587922

**Authors:** Eri Hirata, Ken-taro Sakata, Grace I. Dearden, Faria Noor, Indu Menon, George N. Chiduza, Anant K. Menon

## Abstract

The oligosaccharide needed for protein *N*-glycosylation is assembled on a lipid carrier via a multi-step pathway. Synthesis is initiated on the cytoplasmic face of the endoplasmic reticulum (ER) and completed on the luminal side after transbilayer translocation of a heptasaccharide lipid intermediate. More than 30 Congenital Disorders of Glycosylation (CDGs) are associated with this pathway, including RFT1-CDG which results from defects in the membrane protein Rft1. Rft1 is essential for the viability of yeast and mammalian cells and was proposed as the transporter needed to flip the heptasaccharide lipid intermediate across the ER membrane. However, other studies indicated that Rft1 is not required for heptasaccharide lipid flipping in microsomes or unilamellar vesicles reconstituted with ER membrane proteins, nor is it required for the viability of at least one eukaryote. It is therefore not known what essential role Rft1 plays in *N*-glycosylation. Here, we present a molecular characterization of human Rft1, using yeast cells as a reporter system. We show that it is a multi-spanning membrane protein located in the ER, with its N and C-termini facing the cytoplasm. It is not *N*-glycosylated. The majority of RFT1-CDG mutations map to highly conserved regions of the protein. We identify key residues that are important for Rft1’s ability to support *N*-glycosylation and cell viability. Our results provide a necessary platform for future work on this enigmatic protein.

## INTRODUCTION

Asparagine-linked (*N*-linked) protein glycosylation is found in all three domains of life. In eukaryotes, it occurs in the lumen of the endoplasmic reticulum (ER) where the enzyme oligosaccharyltransferase (OST) attaches a pre-synthesized oligosaccharide (Glucose_3_Mannose_9_*N*-acetylglucosamine_2_ (abbreviated Glc_3_Man_9_GlcNAc_2_, or simply G3M9) in yeast and humans) to asparagine residues within glycosylation sequons in newly translocated proteins (Fig. 1A) (1, 2). The oligosaccharide is assembled by sequentially glycosylating the isoprenoid lipid carrier dolichyl phosphate. This occurs in two stages whereby the lipid intermediate Man_5_GlcNAc_2_-PP-dolichol (M5-DLO) is generated on the cytoplasmic face of the ER, then elaborated to the mature glycolipid G3M9-DLO after being flipped across the membrane to the luminal side (Fig. 1A) (3–8). The molecular identities of the necessary glycosyltransferases are known, and several of these enzymes have been structurally characterized (1). However, the identity of the protein that flips M5-DLO across the ER membrane is controversial — biochemical studies indicate that it is a scramblase-type lipid transporter capable of equilibrating M5-DLO across the membrane, in an ATP-independent manner and with high specificity (older reports refer to this transporter as a (ATP-independent) flippase) (3–6). Defects in *N*-glycosylation underlie numerous human genetic disorders including a heterogeneous group of autosomal-recessive, metabolic diseases termed Congenital Disorders of Glycosylation (CDGs) (9–14). More than 30 CDGs are associated with the core reactions needed to synthesize *N*-glycoproteins in the ER.

**Figure 1.**
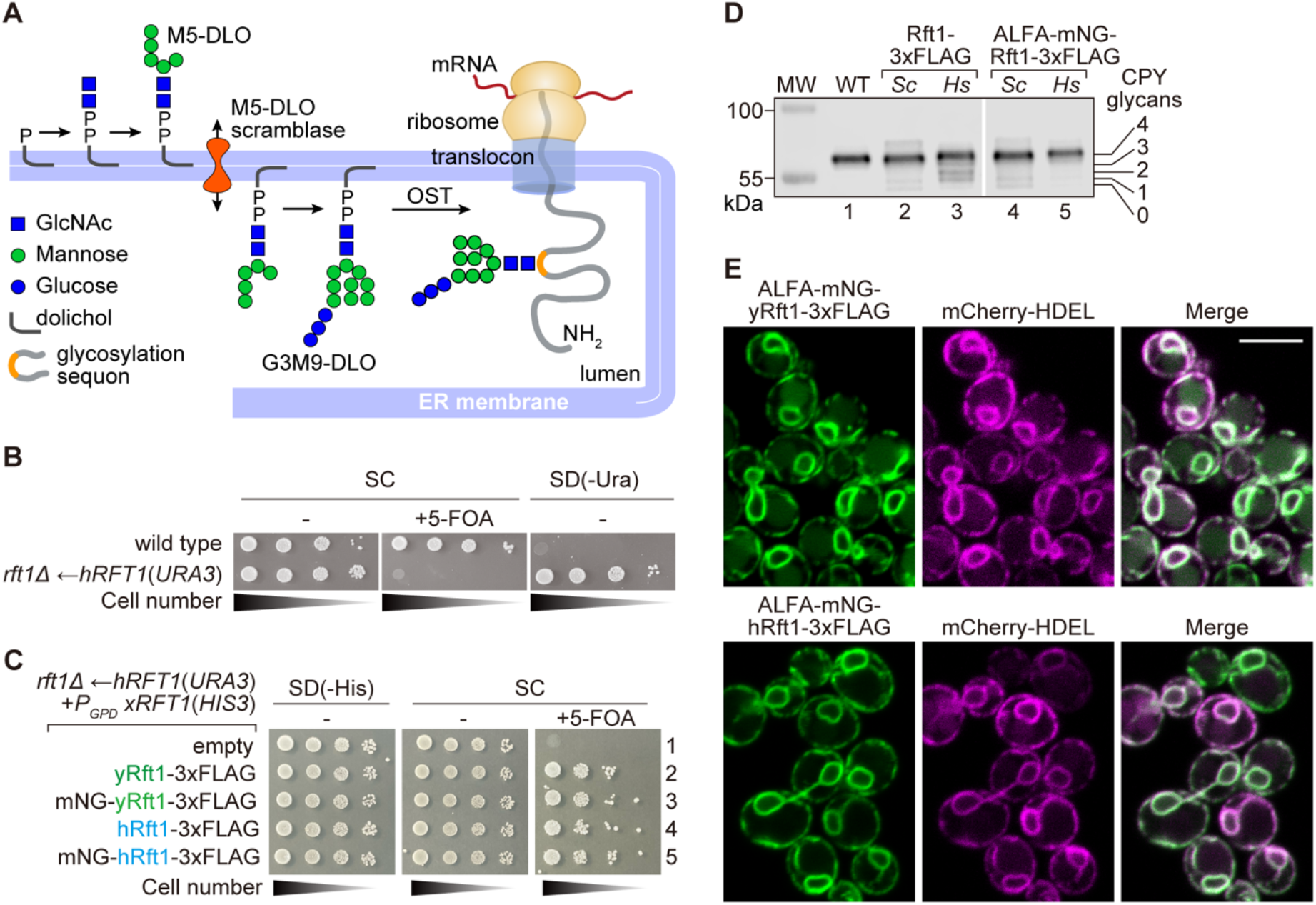
Functional replacement of yeast Rft1 by human Rft1. A. Protein *N*-glycosylation in the ER. Glc_3_Man_9_GlcNAc_2_-PP-dolichol (G3M9-DLO), the oligosaccharide donor for protein *N*-glycosylation in yeast and humans, is synthesized in two stages. The first stage produces Man_5_GlcNAc_2_-PP-dolichol (M5-DLO) on the cytoplasmic side of the ER. M5-DLO then moves across the membrane to the luminal side (a process facilitated by M5-DLO scramblase) where it is converted to G3M9-DLO. Oligosaccharyltransferase (OST) transfers the G3M9 oligosaccharide from G3M9-DLO to a glycosylation sequon (NXS/T, single letter amino acid code, where X is any amino acid except proline) in a nascent protein as it emerges from the protein translocon into the ER lumen. B. Serial 10-fold dilutions of wild-type (BY4741) and KSY512 (*rft1Δ←*hRft1(*URA3*)) cells were spotted on SC plates +/- 5-FOA and SD(-Ura) and incubated at 30°C for 3 days. C. KSY512 (*rft1Δ←*hRft1(*URA3*)) cells were transformed with an empty *HIS3* vector (empty, p413-*P_GPD_*), or *HIS3* vectors encoding various Rft1 constructs (xRft1) as follows: yRft1-3xFLAG, hRft1-3xFLAG, ALFA-mNG-yRft1-3xFLAG, and ALFA-mNG-hRft1-3xFLAG. The cells were spotted (10-fold serial dilutions) on the indicated media and photographed after incubation at 30°C for 3 days. D. Wild-type (BY4741) and KSY512 cells transformed with various Rft1 constructs as in panel C, were cultured in YPD liquid medium to log-phase, harvested and analyzed by SDS-PAGE and immunoblotting with anti-CPY antibody. The annotation on the right indicates migration of mature CPY (with 4 N-glycans), and hypoglycosylated forms with 3, 2, 1 or 0 glycans. E. Fluorescently-tagged mNG-yRft1 and mNG-hRft1 constructs (expression driven by the GPD promoter) were integrated into wild-type cells expressing the luminal ER marker mCherry-HDEL. The resulting YAKM301 (P_GPD_-mNG-yRft1 mCherry-HDEL) and YAKM225 (P_GPD_-mNG-hRft1 mCherry-HDEL) cells were cultured in YPD medium to log-phase and imaged by confocal fluorescence microscopy. Scale bar = 5 μm.

RFT1-CDG is associated with defects in the membrane protein Rft1 (15–17), which was proposed as the M5-DLO scramblase based on genetic studies in yeast (18–20). Rft1 is essential for the viability of yeast (19) and mammalian cells (https://depmap.org/portal/), and its deficiency results in the accumulation of M5-DLO, depletion of G3M9-DLO and hypoglycosylation of *N*-glycoproteins (18, 19). This happens even though the enzymes that convert M5-DLO to G3M9-DLO are intact and OST is unaffected. Importantly, although OST can use M5-DLO as a glycan donor (1), this does not appear to occur in Rft1-depleted cells. One explanation of these data is that in the absence of the purported Rft1 scramblase, M5-DLO accumulates on the cytoplasmic face of the ER where it cannot be used for G3M9-DLO synthesis and/or the OST reaction. However, where M5-DLO accumulates in Rft1-deficient cells is not known.

In contrast to these suggestive results pointing to a role for Rft1 in scrambling M5-DLO in cells, biochemical studies indicated that Rft1 is not necessary for scrambling *in vitro*. Thus, we recapitulated M5-DLO scrambling in large unilamellar vesicles reconstituted with a “Triton Extract (TE)”, a mixture of all ER membrane proteins selectively extracted with detergent from yeast or rat liver microsomes (5, 6, 21, 22). We found that scrambling in this system is very specific, with higher order glycolipids (Glc_1-2_Man_6-9_GlcNAc_2_-PP-dolichol) and a structural isomer of M5-DLO being scrambled at least an order of magnitude more slowly than M5-DLO (5, 6). Quantitative elimination of Rft1 from the TE by affinity chromatography prior to reconstitution did not impact scrambling, indicating that Rft1 is dispensable for activity. In other experiments, we resolved TE proteins by different methods including dye-resin chromatography and velocity gradient sedimentation. When fractions from these separations were reconstituted into liposomes on an equivalents’ basis and assayed, M5-DLO scramblase activity could be clearly resolved from Rft1 (5, 22). These results indicate that even if Rft1 has scramblase activity, it’s contribution to the overall M5-DLO scramblase activity of the TE is minor.

In a different study, microsomes were prepared from yeast cells in which Rft1 levels had been lowered by using a regulatable promoter to drive down protein expression (23). Analysis of the cells showed the expected phenotype of Rft1 deficiency, i.e., accumulation of M5-DLO and hypoglycosylation of the reporter glycoprotein carboxypeptidase Y (CPY). However, intact microsomes derived from these cells were able to execute the entire DLO biosynthetic sequence, converting newly synthesized GlcNAc_2_-PP-dolichol to Man_9_GlcNAc_2_-PP-dolichol, implying that they have M5-DLO scramblase activity. This was confirmed by direct measurement of scrambling using the M5-DLO analog GlcNAc_2_-PP-dolichol_15_. These results suggest that the inability of Rft1-deficient cells to use M5-DLO is not due to the absence of scramblase activity but possibly because some feature of ER architecture in intact cells poses a barrier that is lost when the cells are disrupted.

An essential role for Rft1 in scrambling M5-DLO was also questioned in a study of the protein in the early diverging eukaryote *Trypanosoma brucei* (24). Rft1-null procyclic (insect-stage) trypanosomes were found to grow normally. They had normal steady state levels of mature DLO and significant *N*-glycosylation consistent with sufficient M5-DLO scramblase activity, yet accumulated M5-DLO to a steady state level 30-100-fold greater than found in wild-type cells.

The cumulative data presented above pose a conundrum in that Rft1 is not required for M5-DLO scramblase activity *in vitro* (reconstituted vesicles and microsomes) or in *T. brucei* cells, but clearly plays a role in the metabolic fate of M5-DLO and is essential for the viability of yeast and human cells. Thus, it remains to be determined what essential role Rft1 might play in the cell and how its deficiency results in RFT1-CDG. As a first step towards resolving these issues, we chose to characterize the protein. Rft1 is relatively under-studied, with no published reports concerning its sub-cellular localization, membrane topology and structure-function aspects in yeast and human cells. Here, we present a molecular characterization of human Rft1 (hRft1), using the yeast *Saccharomyces cerevisiae*, which is well-established as a model system for the study of CDGs (25). We show that human Rft1 functionally substitutes for its yeast counterpart. It is localized throughout the ER and has a polytopic arrangement with its N and C-termini facing the cytoplasm. The majority of RFT1-CDG mutations map to highly conserved regions of the protein. Using a suite of assays to correlate protein expression, CPY *N*-glycosylation and cell doubling time, we identify key residues that are important for Rft1’s ability to support glycosylation and cell viability. These results provide a necessary platform for future work on Rft1.

## RESULTS AND DISCUSSION

### Functional replacement of yeast Rft1 by human Rft1

We transformed a heterozygotic diploid yeast strain (*rft1::KANMX4 / RFT1*) (26) with a *URA3* plasmid (p416-*P_GPD_*, hRft1-3xFLAG or simply hRft1(*URA3*)) (Table 1) encoding human Rft1 (hRft1) with a C-terminal 3xFLAG tag. The transformed cells were sporulated, and individual spores lacking endogenous *RFT1* but carrying the hRft1-expressing *URA3* plasmid were selected on the basis of G418 resistance and ability to grow on plates lacking uracil. We chose cells (henceforth termed KSY512) with mating type ’a’ for subsequent experiments.

**Table 1.**
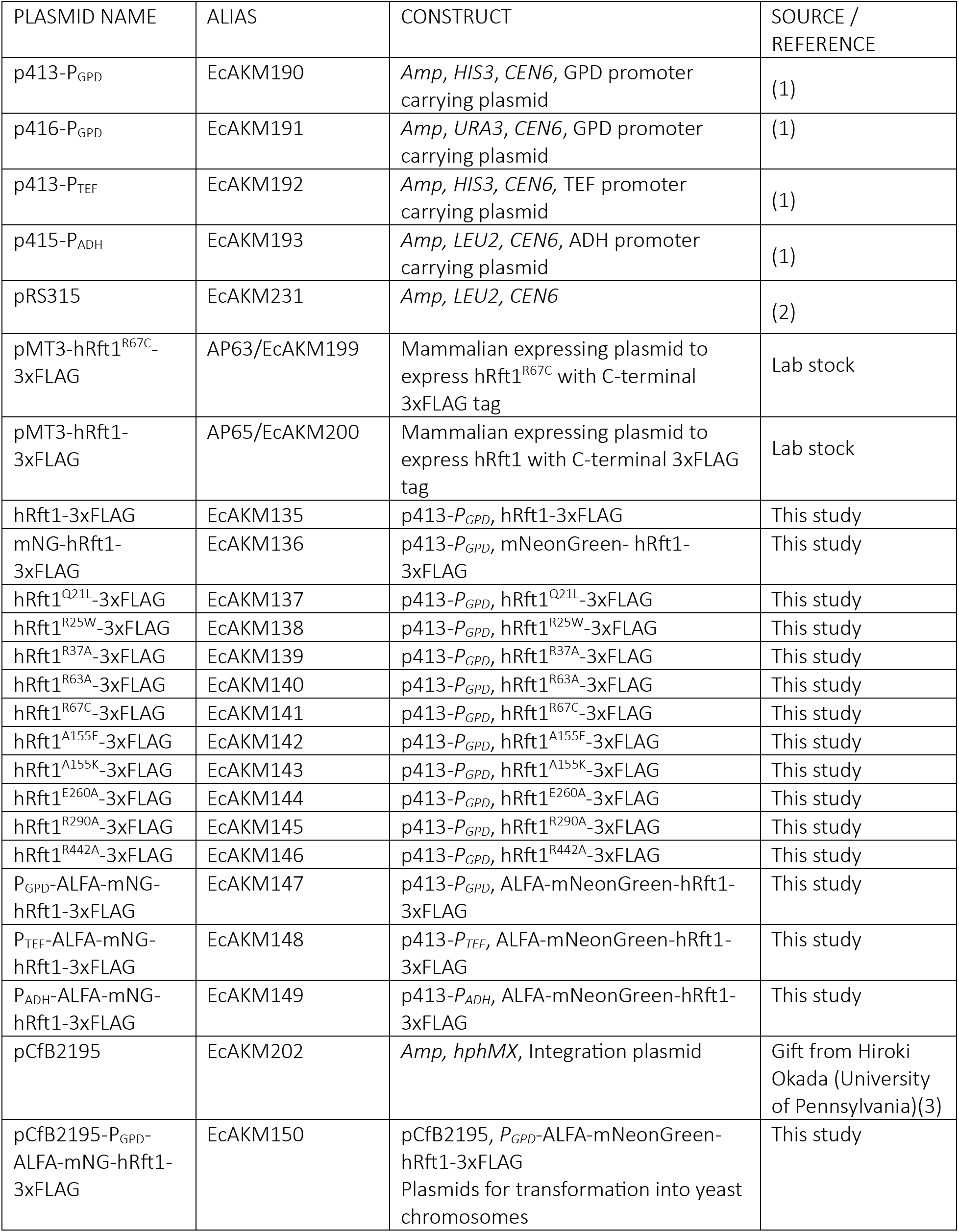
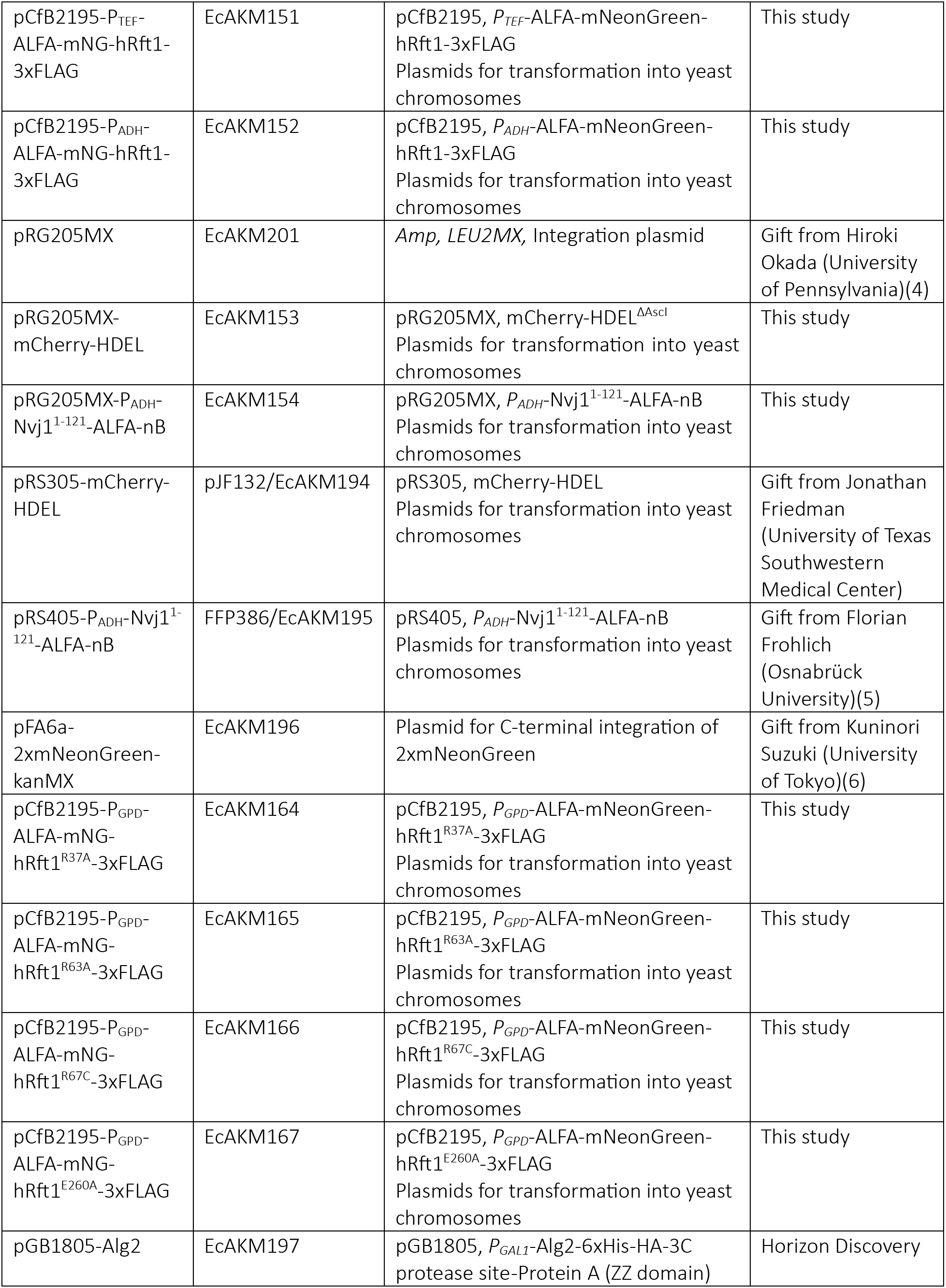
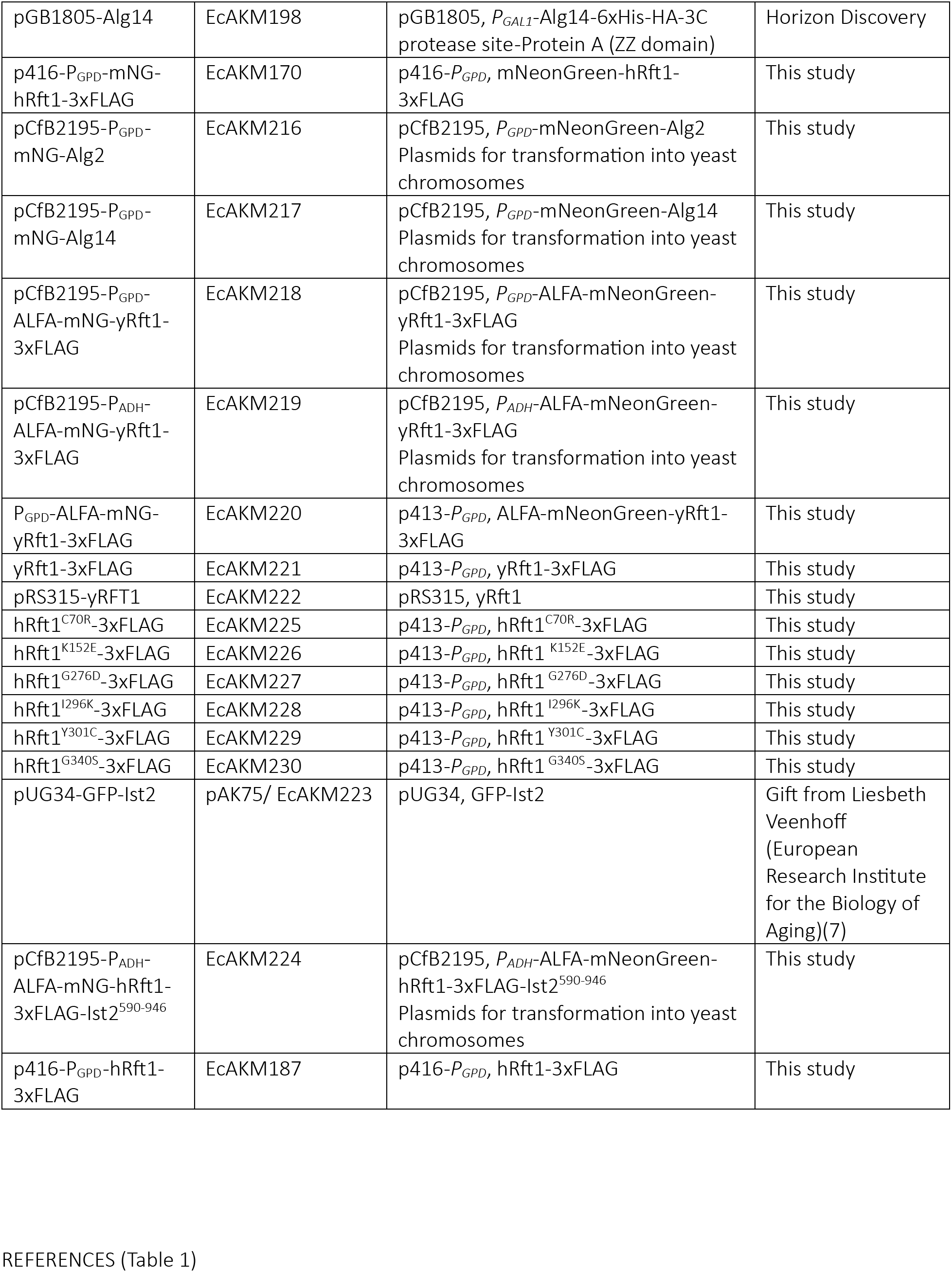
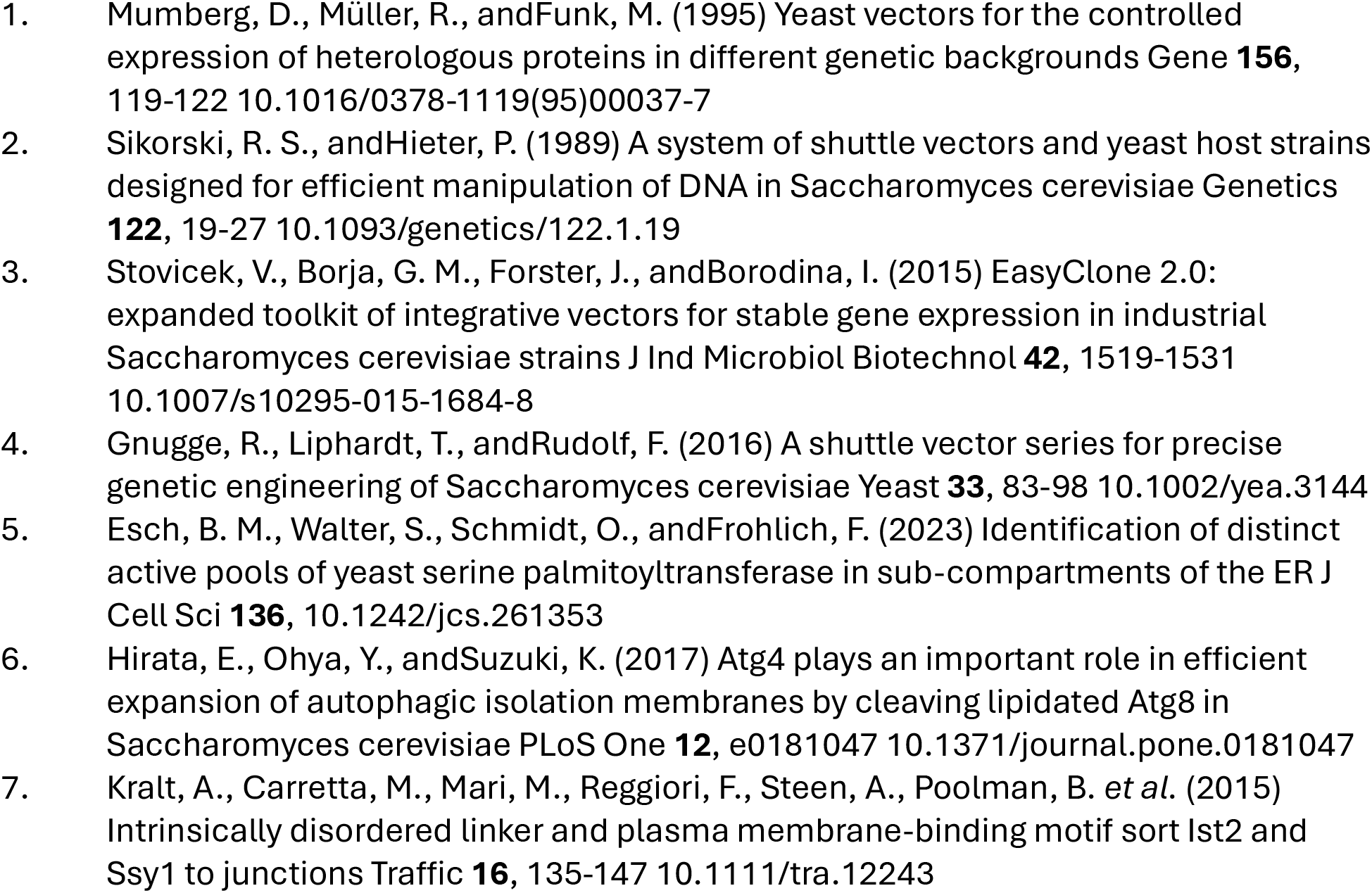
Plasmids.

The KSY512 cells (*rft1Δ←*hRft1(*URA3*)) grew on synthetic Ura-media (SD(-Ura)) as expected but did not grow on plates containing 5-fluoroorotic acid (5-FOA) (Fig. 1B), which is converted by the plasmid-encoded *URA3* gene product into toxic 5-fluorouracil. This indicates that the hRft1(*URA3*) plasmid is essential for the viability of KSY512 cells. In contrast, isogenic wild-type BY4741 cells did not grow on Ura-media but grew on 5-FOA (Fig. 1B). We conclude that yeast Rft1 (yRft1) can be functionally replaced by hRft1, as noted previously (15), and that the C-terminal 3xFLAG tag does not appear to affect hRft1 function significantly (but see below), consistent with a previous report in which functional yRft1 was expressed with a C-terminal Protein A tag (21).

To quantify the functionality of hRft1-3xFLAG in the yeast system we compared it with yRft1-3xFLAG. Thus, we transformed KSY512 cells with *HIS3* plasmids encoding hRft1-3xFLAG or yRft1-3xFLAG under control of the constitutively active GPD promoter (27) and tested the cells for growth on 5-FOA plates. In this condition, the hRft1(*URA3*) plasmid in the KSY512 cells will be lost and cell growth will depend on the Rft1 variants expressed from the *HIS3* plasmid, enabling a direct comparison of their function. Figure 1C (right panel, compare rows 2 and 4), shows that both proteins supported growth. We picked colonies from the 5-FOA plate (corresponding to *rft1Δ←*hRft1(*HIS3*) and *rft1Δ←*yRft1(*HIS3*) cells) to analyze the steady-state *N*-glycosylation status of CPY, a vacuolar protein with four *N*-glycans (28). Protein extracts from the cells were processed for SDS-PAGE followed by immunoblotting with anti-CPY antibodies, and the resulting pattern of CPY glycoforms was quantified to obtain a glycosylation score (Glycoscore) as previously described (29). The CPY profile in wild-type cells as well as yRft1 and hRft1-expressing cells was dominated by a single band corresponding to fully *N*-glycosylated mature CPY (Fig. 1D, lanes 1-3), with some hypoglycosylation in the latter cells as evinced by a faint ladder of lower molecular weight bands corresponding to CPY with <4 *N*-glycans (Fig. 1D, lanes 2 and 3). The corresponding Glycoscores (mean ± S.D. (n=3)) were 93.4 ± 1.4 (wild type) versus 83.6 ± 2.4 (yRft1) and 77.2 ± 2.4 (hRft1). The hierarchy of CPY Glycoscores in the three samples (wild type > yRft1 > hRft1) may be explained by the presence of the 3xFLAG tag and/or to differences in expression level of the yRft1 and hRft1 proteins. We compared the expression of hRft1-3xFLAG and yRft1-3xFLAG by SDS-PAGE and immunoblotting with anti-FLAG antibodies. Figure S1A (lanes 1 and 2) shows that both proteins migrate faster than expected based on their predicted molecular weights, likely due to detergent binding effects which are known to cause anomalous migration of membrane proteins in SDS-PAGE analyses (30), with yRft1-3xFLAG being expressed at a ∼7-fold higher level. The reason for the difference in expression level is not clear. Quantitative immunoblotting in comparison to a 3xFLAG protein standard revealed that hRft1-3xFLAG is expressed at about 800 copies per cell (Fig. S1B, C), comparable to the reported copy number for endogenous yeast Rft1 (∼1000 copies/cell (31)); in contrast plasmid-expressed yRft1-3xFLAG is produced at ∼5500 copies/cell. The presence of the 3xFLAG-tag in both proteins together with the difference in expression level likely explains the observed differences in their ability to support CPY glycosylation compared with wild-type cells.

### Rft1 localizes to the ER

We next examined the sub-cellular localization of Rft1. For this, we generated fluorescently tagged hRft1 and yRft1 constructs, with mNeonGreen (mNG) (32), a monomeric green/yellow fluorescent protein, fused to its N-terminus. The construct also contains an ALFA-tag (33), N-terminal to mNG, and a C-terminal 3xFLAG tag.

We first tested whether the mNG-tagged constructs are functional. We therefore transformed KSY512 cells with *HIS3* plasmids encoding the tagged proteins under control of the GPD promoter, then tested the cells for growth on 5-FOA plates. Figure 1C (right panel, rows 3 and 5) show that both constructs are functional as the cells grow on 5-FOA. We picked colonies from the 5-FOA plate to analyze expression level of the constructs and quantify CPY glycosylation. We found that both constructs are comparably expressed (Fig. S1C (lanes 3 and 4)) and able to support CPY glycosylation (Fig. 1D, lanes 4 and 5) yielding CPY Glycoscores (mean ± S.D. (n=3)) of 83.8 ± 4.1 (mNG-yRft1) and 85.6 ± 1.7 (mNG-hRft1). Of note, the N-terminal tag appeared to have a stabilizing effect on hRft1, making its expression comparable to that of the more highly expressed yeast protein, with an associated improvement in CPY glycosylation score.

To investigate the sub-cellular localization of the proteins by fluorescence microscopy we integrated the constructs in the genome of wild-type cells expressing the luminal ER marker mCherry-HDEL. Figure 1E shows that both proteins display a characteristic yeast ER pattern, mainly comprising cortical and nuclear ER regions and overlapping precisely with the distribution of mCherry-HDEL. As this pattern was also observed for mNG-hRft1 constructs expressed under the control of constitutive promoters of different strengths (GPD>TEF>ADH) (Fig. S2) (27) it is not the result of mis-localization due to saturation of trafficking mechanisms. Thus, hRft1 and yRft1 are ER-localized proteins.

### Is Rft1 localized to an ER domain?

We previously proposed that early steps of DLO synthesis may be laterally segregated from the M5-DLO scramblase in the ER (5), and that a possible role of Rp1 may be to chaperone M5-DLO within the plane of the membrane, from its site of synthesis to the scramblase. This would account for why Rp1 is important in cells where DLO synthesis is laterally compartmentalized, but not in reconstituted systems or in microsomes where compartmentalization is lost. A related scenario was proposed for Lec35/MPDU1, another protein with an enigmatic role in *N*-glycosylation (34, 35).

Consistent with our idea, Krahmer et al. (36) reported that several enzymes needed to convert dolichyl-P to M5-DLO on the cytoplasmic face of the ER (Fig. 1A) — including Alg14, the membrane-bound subunit of the heterodimeric GlcNAc transferase complex (37–39), and the mannosyltransferases Alg1, Alg2 and Alg11 — can be co-isolated with lipid droplets (LDs) in oleic acid-fed insect cells. They also reported that Rft1 was located exclusively in the LD fraction. In contrast, the luminally-oriented glycosyltransferases that convert M5-DLO to G3M9-DLO do not associate with LDs. The hydrophobic portion of Alg14 adopts conformations that enable it to integrate into membrane bilayers as well as phospholipid monolayers that surround LDs (40, 41), accounting for its enrichment in the LD proteome. The mechanism by which Alg1, Alg2, Alg11 and Rft1 might associate with LDs is likely different — these membrane proteins may reside in a domain of the ER that wraps around LDs (42) and thus co-isolates with these organelles. These observations suggest that the DLO pathway is not only transversely segregated across the ER membrane as depicted in Figure 1A, but also laterally compartmentalized with M5-DLO being generated in an ER domain that can be isolated and visualized via its association with LDs.

To test this scenario in our yeast model, we integrated mNG-tagged Alg14 and Alg2 into the genome of wild-type yeast cells expressing the LD marker Erg6-mCherry (43). The cells were grown in rich medium supplemented with oleic acid to induce LDs and examined by fluorescence microscopy. We found that Alg14 co-localized strongly with LDs, visualized as ring-like structures marked by Erg6-mCherry (Fig. 2A), whereas Alg2 displayed a typical ER pattern, distinct from LDs (Fig. 2B). We next integrated mNG-yRft1 and mNG-hRft1 in the genome of the Erg6-mCherry-expressing cells and examined their localization after LD induction. Figure 2 (panels C and D) show that the mNG-Rft1 proteins retain their pan-ER distribution (as in Fig. 1E and Fig. S2) in oleic acid-fed cells distinct from the Erg6-mCherry-marked LD structures. We conclude that the synthesis of M5-DLO is partly spatially restricted as evinced by co-localization of Alg14 with LDs. However, the Alg2 mannosyltransferase and Rft1 do not colocalize with LDs as reported previously (36).

**Figure 2.**
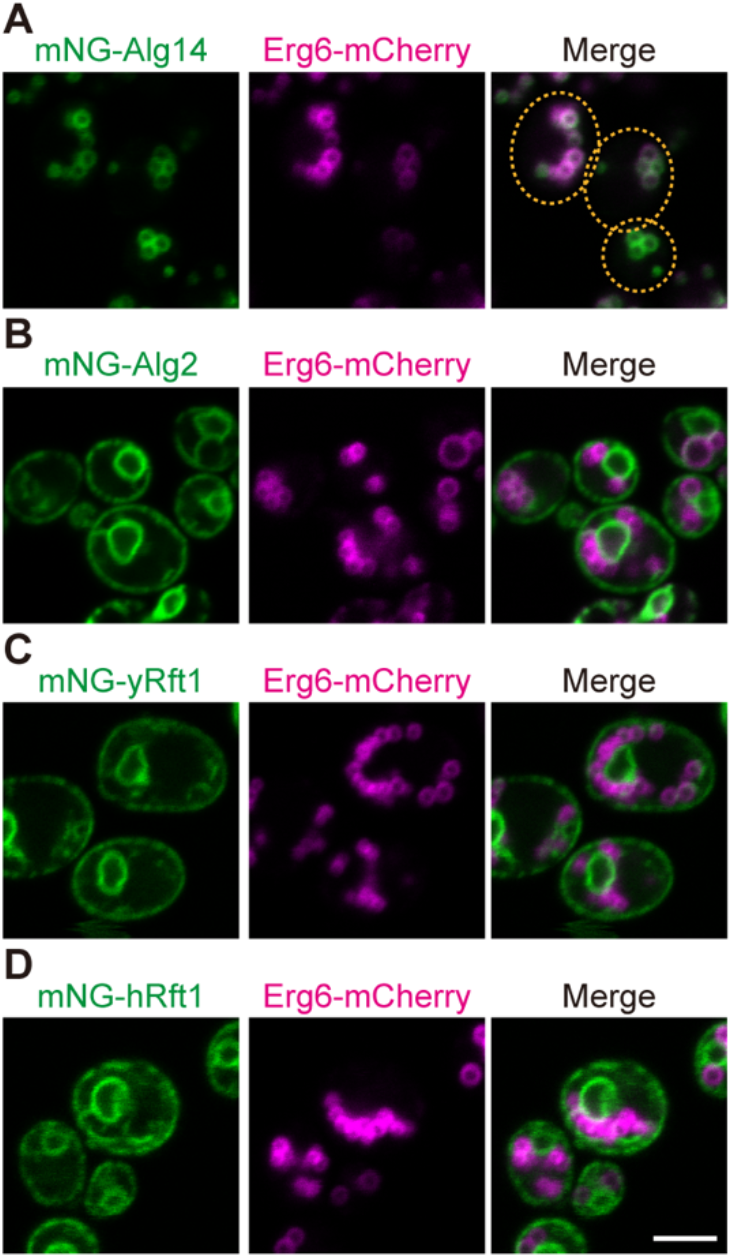
Alg14 co-localizes with lipid droplets, but Alg2 and Rft1 do not. A. YAKM275 (P_GPD_-mNG-Alg14 Erg6-mCherry) cells expressing fluorescently tagged Alg14 and Erg6 were cultured in medium supplemented with oleic acid (YPO medium) for 16 h and visualized by confocal fluorescence microscopy. The dotted line indicates the shape of the cells. B. As in panel A, except that YAKM274 (P_GPD_-mNG-Alg2 Erg6-mCherry) cells expressing fluorescently tagged Alg2 and Erg6 were visualized. C-D. As in panel A, except that YAKM302 (P_ADH_-mNG-yRft1 Erg6-mCherry) and YAKM235 (P_ADH_-mNG-hRft1 Erg6-mCherry) cells were visualized. Scale bar = 5 μm for all panels.

### Rft1 is not necessary for M5-DLO scramblase activity in liposomes reconstituted with total ER membrane proteins

We next re-assessed the hypothesis that Rft1 is responsible for scrambling M5-DLO across the ER membrane. We prepared a salt-washed microsomal fraction from a homogenate of KSY512 (*rft1Δ←*hRft1(*URA3*)) cells, selectively extracted ER membrane proteins with ice-cold Triton X-100 as previously described (5) and reconstituted the protein mixture (Triton Extract (TE)) into large unilamellar vesicles composed of egg phosphatidylcholine (egg PC), with trace quantities of [^3^H]M5-DLO and a fluorescent PC analog, NBD-PC, which has a nitrobenzoxadiazole (NBD) fluorophore attached to one of its acyl chains. In parallel, we incubated an identical sample of TE with anti-FLAG resin to remove hRft1-3xFLAG, before reconstituting it into LUVs. Protein-free liposomes were also prepared. Immunoblotting analysis showed that anti-FLAG treatment quantitatively and specifically removed hRft1-3xFLAG from TE — no FLAG signal was detected in the treated sample, whereas the signal corresponding to an irrelevant ER protein, Dpm1, was unchanged (Fig. 3A). Dynamic light scattering measurements indicated that the vesicle samples were similar, with an average diameter of ∼175 nm (Fig. 3B).

**Figure 3.**
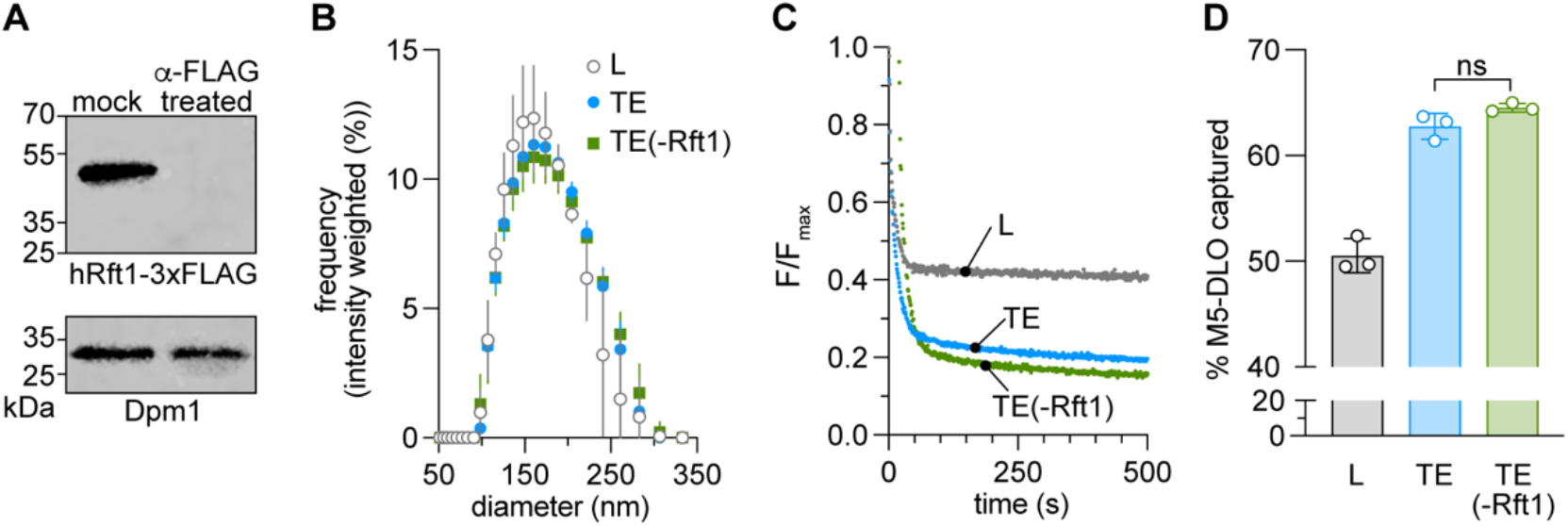
Rft1 is not necessary for M5-DLO scrambling in vesicles reconstituted with yeast ER membrane proteins. Microsomes were prepared by differential centrifugation of a homogenate of KSY512 cells, and salt-washed to remove peripheral proteins. The salt-washed membranes were extracted with ice-cold Triton X-100 to solubilize ER membrane proteins. The Triton extract (TE) was mock-treated or incubated with anti-FLAG resin to eliminate hRft1-3xFLAG, then reconstituted with egg phosphatidylcholine and trace quantities of NBD-PC and [^3^H]M5-DLO to generate large unilamellar proteoliposomes (indicated as ’TE’ and ’TE(-Rft1)’) for scramblase activity assays. The protein/phospholipid ratio of the proteoliposomes was ∼45 mg/mmol, based on input values of protein and phospholipid. Protein-free liposomes (L) were prepared in parallel. A. Immunoblot using anti-FLAG (top) and anti-Dpm1 (bottom) antibodies. Identical cell equivalents were loaded in the mock-treated and anti-FLAG resin-treated samples. No FLAG signal was detected in the anti-FLAG resin-treated sample even upon loading 10-times more sample (not shown). B. Diameter of reconstituted vesicles measured by dynamic light scattering. Error bars=mean ± S.D. (n=3 technical replicates). C. NBD-PC scramblase activity assay. Dithionite was added at t=0 s and fluorescence (F) was monitored over time. The TE and TE(-Rft1) traces (F/F_max_, normalized to the average fluorescence (F_max_) prior to dithionite addition) overlap exactly; to improve visualization, the TE(-Rft1) trace is displaced downwards (0.05 y-units) and to the right (20 x-units). D. M5-DLO scramblase activity assay. The y-axis indicates the fraction of [^3^H]M5-DLO in the reconstituted vesicles that is captured by exogenously added Con A. For liposomes, the % capture is predicted to be 50%; for proteoliposomes with M5-DLO scramblase activity, the capture efficiency increases, the exact amount depending on the fraction of vesicles that has scramblase activity. See text for details. Error bars=mean ± S.D. (n=3 technical replicates); ns, no significant difference using ordinary one-way ANOVA.

We previously showed that proteoliposomes reconstituted with yeast TE have glycerophospholipid scramblase activity (5, 44), reflecting the biogenic properties of the ER (3, 4, 45). As this activity is not expected to be related to Rft1, we used it as a quality control test to compare proteoliposomes prepared with anti-FLAG-resin-treated TE versus mock-treated TE. We assayed the activity as previously described by measuring the ability of the membrane-impermeant reductant dithionite to bleach NBD-PC molecules in the outer leaflet of reconstituted vesicles. For protein-free liposomes, NBD-PC molecules in the inner leaflet are protected, whereas for proteoliposomes containing scramblases, these molecules are scrambled to the outer leaflet where they are exposed to dithionite. Thus, approximately half the fluorescence of protein-free liposomes is expected to be lost on dithionite treatment, whereas a larger proportion of fluorescence should be lost in a vesicle population where some or most vesicles contain a functional scramblase. This is indeed what we observed (Fig. 3C) — dithionite treatment of protein-free liposomes resulted in a drop of ∼57% of fluorescence, whereas for proteoliposomes, irrespective of whether they were generated from anti-FLAG-resin-treated TE or mock-treated TE, the drop was ∼80% (the traces are identical, hence they are shown displaced from one another for easy visualization (Fig. 3C)). These numbers indicate that, at the protein/phospholipid ratio used for reconstitution, approximately 55% of the vesicles contain a phospholipid scramblase (calculated as described (46)). Furthermore, the identical fluorescence reduction traces obtained for proteoliposomes generated from the treated and mock-treated samples indicate that these vesicles are similarly reconstituted.

We next assayed the same vesicles for M5-DLO scramblase activity, using a previously described assay (5, 21, 22) in which the mannose-binding lectin Concanavalin A (Con A) is used to capture M5-DLO molecules located in the outer leaflet of the vesicles. For protein-free vesicles, approximately half the M5-DLO molecules are expected to be captured at the endpoint of the assay as the remainder are confined to the inner leaflet. For proteoliposomes with M5-DLO- scramblase activity, all M5-DLO molecules are expected to be captured as those from the inner leaflet are translocated to the Con A-accessible outer leaflet. As shown in Figure 3D, ∼50% of M5- DLO is captured in protein-free liposomes as expected, whereas ∼65% is captured in both types of proteoliposomes. Thus, the presence or absence of Rft1 does not affect the outcome of the M5- DLO scramblase assay. We note that a larger proportion of proteoliposomes contain phospholipid scramblase activity compared with M5-DLO scramblase activity, ∼55% versus ∼30%, respectively (calculated as described (46)), consistent with the greater abundance of phospholipid scramblase(s) in the TE as reported previously (5).

Thus, proteoliposomes generated from an extract of ER membrane proteins, *sans* Rft1, have undiminished M5-DLO scramblase activity compared with Rft1-replete vesicles. This updated result, building on a new yeast test strain expressing hRft1 and incorporating additional quality controls for vesicle reconstitution, extends our previous conclusion (21) that Rft1 is not required for scrambling in our reconstitution-based assay. If Rft1 does indeed have M5-DLO scramblase activity, a possibility that can be tested in the future when high-quality purified protein is available, it would appear to be a minor contributor, and redundant with other scramblases as our reconstitution data clearly indicate that the majority of the scramblase activity in the TE is due to another protein(s). A back-of-the-envelope analysis of vesicle occupancy data determined in a previous reconstitution-based study of M5-DLO scramblase activity (5) indicates — with standard assumptions concerning vesicle diameter, cross-sectional area of a phospholipid and the average molecular mass of ER membrane proteins — that the M5-DLO scramblase represents ≥ 1% by weight of proteins in the TE. As there are approximately 2 x 10^6^ ER membrane proteins in a haploid yeast cell (47), we conclude that there are at least 20,000 M5-DLO scramblases per cell. In contrast, there are only <1000 copies of hRft1 per cell in the KSY512 strain used for the present analysis (Fig. S1B, C).

### Architecture of hRft1

Structure prediction by AlphaFold2 (48, 49) reveals that hRft1 has 14 transmembrane (TM) spans (Fig. 4A). This model is of high quality as judged by the predicted Local Distance Difference Test (pLDDT) (48) score which measures local confidence in the structure on a per residue basis. Scores range from 0–100, with >90 representing high accuracy and >70 corresponding to correct backbone prediction. Although residues in a portion of TM1, the disordered C terminus, and intracellular loop 3 (ICL3) had predicted pLDDT scores below 70, the remaining residues had higher scores (70–98.56) indicating the overall good quality of the model. Of note, whereas the lower pLDDT scores impact the quality of the structure predictions in certain regions of the protein, they do not affect the overall 14-TM topology prediction. We also constructed an hRft1 model with DeepTMHMM (50), a program which uses a pretrained protein language model to predict membrane protein topology, as well as orientation in the membrane. For hRft1, DeepTMHMM indicates a 14-TM model similar to that produced by AlphaFold2, with the additional prediction that the N-terminus and C-terminus of the protein are located on the cytoplasmic side of the membrane.

**Figure 4.**
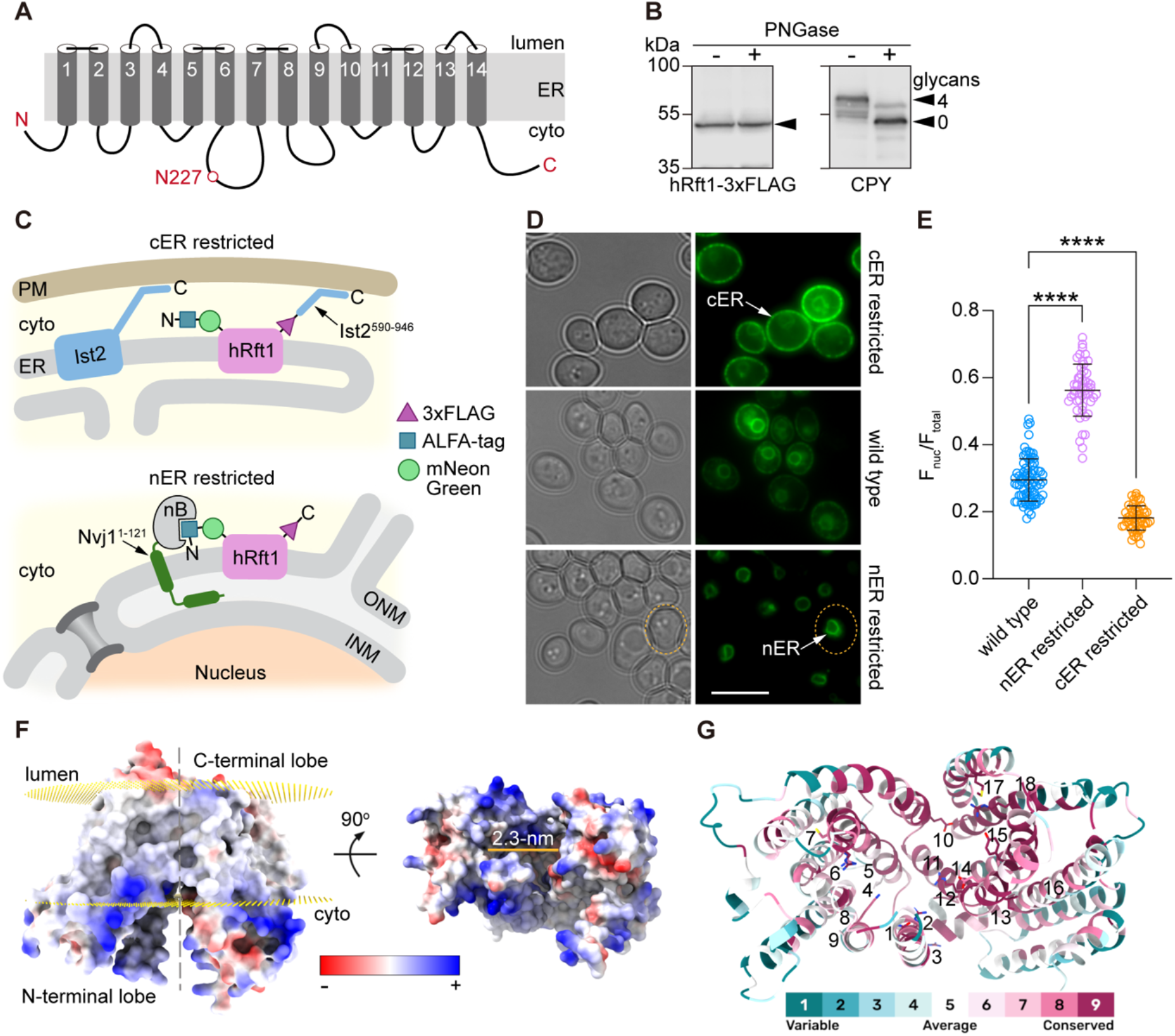
Functional architecture of Rft1. A. Topology model of hRft1. The model is based on DeepTMHMM (https://dtu.biolib.com/DeepTMHMM) which predicts 14 transmembrane spans. The protein has its N- and C-terminus oriented towards the cytoplasm. The only *N*-glycosylation sequon (N^227^IT) is located in the third intracellular loop (ICL3). The relative lengths of the loops are shown roughly to scale. B. hRft1 is not *N*-glycosylated. A protein extract from hRft1-3xFLAG-expressing KSY512 cells was treated with PNGase F (a control sample was mock-treated in parallel) and subsequently analyzed by SDS-PAGE immunoblotting using anti-FLAG antibodies (to detect hRft1) and anti-CPY antibodies. Left panel, arrowhead indicates migration of hRft1. Right panel, arrowheads indicate the positions of fully glycosylated (4 glycans) and non-glycosylated (0 glycans) CPY; tick marks represent the same molecular weight markers as shown in the left panel. C. Fluorescence microscopy assay to test the N_in_, C_in_ orientation of hRft1 in the ER membrane. Top panel, the C-terminal domain of Ist2 (residues 590-946) which contains a plasma membrane binding domain is fused to the C-terminus of ALFA-mNG-hRft1-3xFLAG. When expressed in yeast, the fusion protein is expected to be enriched in the cER. Bottom panel, C-terminally ALFA-nB-tagged Nvj1^1-121^ is expressed together with P_ADH_-ALFA-mNG-hRft1 in yeast cells. As the ALFA-nB tag binds to the N-terminal ALFA tag of hRft1, the protein is expected to be enriched in the nER. D. YAKM172 (P_tet_-Rft1 P_ADH_-ALFA-mNG-hRft1), YAKM173 (P_tet_-Rft1 Nvj1-nB P_ADH_-ALFA-mNG-hRft1), and YAKM287 (P_tet_-Rft1 P_ADH_-ALFA-mNG-hRft1-Ist2^590-946^) were visualized by wide-field microscopy (brightfield, left panels; fluorescence, right panels). The middle panels show the normal distribution of ALFA-mNG- hRft1-3xFLAG in cells, similar to images shown in Fig. 1E. Arrows indicate the cER or nER. The dotted line (bottom panel) indicates the shape of an exemplary cell. Scale bar = 5 μm. E. Fluorescence images similar to those shown in panel D were quantified. The total fluorescence (F_total_) and nuclear fluorescence (F_nuc_) of each cell was determined by using ImageJ to measure the fluorescence within approximately circular outlines of the cell and the nucleus. Similar outlines in a cell-free area of the image were used to determine background correction. The graph shows F_nuc_/F_total_ (error bars = mean ± S.D. (n>50)) for wild-type, nER-restricted and cER-restricted samples. ****, p<0.0001 using ordinary one-way ANOVA. F. Isosurface of the AlphaFold model of hRFT1 colored by electrostatics. Position of the lipid bilayer is shown as golden lattices on the left with the luminal side on top and the cytosolic side below as indicated. The structure can be divided (grey dashed line) into two lobes each containing 7 of the 14 TM. The width of the hydrophilic cavity between the lobes is ∼23 Å as measured along the indicated gold line in the right view. G. Cytosolic view of the hRFT1 model colored by ConSurf grade, with higher values indicating greater conservation as indicated in the color bar. CDG-1N associated residues (Table 3) and residues mutated and analyzed in this study are indicated by the numbers (1=Q21, 2=R25, 3=R37, 4=I43, 5=R63, 6=R67, 7=C70, 8=K152, 9=A155, 10=E260, 11=G276, 12=R290, 13=I296, 14=E298, 15=Y301, 16=G340, 17=M408, 18=R442).

We tested key features of the hRft1 topology model depicted in Figure 4A. The model predicts that the only *N*-glycosylation sequon in the protein (N^227^IT) is located in ICL3 where it cannot be glycosylated. To test this, we treated an extract of KSY512 cells with the amidase PNGase F. SDS-PAGE immunoblotting revealed that the quadruply glycosylated CPY glycoform collapsed to a lower molecular weight, non-glycosylated band upon treatment (Fig. 4B, right panel), whereas hRft1 (detected via its C-terminal 3xFLAG tag) was not affected (Fig. 4B, left panel). This result is consistent with the location of N227 in a cytoplasmic loop. Of note, we previously showed that M5-DLO scramblase activity is associated with ER proteins or protein complexes that bind Con A-Sepharose (51) — this point distinguishes Rft1, which we now show is not a glycoprotein, from the protein (or proteins) that contribute the majority of M5-DLO scramblase activity in reconstituted systems, thus supporting the result shown in Figure 3.

Figure 4A indicates that the N- and C-termini of hRft1 are oriented to the cytoplasm, i.e., hRft1 has a N_in_, C_in_ topology. If this is the case, we reasoned that it should be possible to use cytoplasmic tethers, appended to these termini, to recruit the protein to either the nuclear ER (nER) or cortical ER (cER). As the nER and cER are readily distinguishable by fluorescence microscopy, this assay provides an easy read-out. We took advantage of a recently reported nanobody (nB)-based system (52) to restrict localization of an ER membrane protein to the nER. Accordingly, we used our mNG- hRft1 construct which has an ALFA-tag at its N-terminus and expressed it in cells in which an anti-ALFA nB is fused to the membrane anchor of nER-localized Nvj1 (Fig. 4C, bottom). We also made use of a tethering system based on Ist2, an ER-localized, ER-plasma membrane tethering protein (53, 54). The cytoplasmic C-terminal tail of Ist2 (residues 590-946), which consists of an intrinsically disordered linker region terminating in a bimodal cortical sorting sequence (amphipathic helix + basic cluster) for plasma membrane binding, can be used to recruit pan-ER membrane proteins to the cER (55). We attached the Ist2 tail to the C-terminus of mNG-hRft1 to (Fig. 4C, top). As shown in Figure 4D we could recruit ALFA-mNG-hRft1-3xFLAG to the nER using the nB-Nvj1-based system (Fig. 4D, bottom), and ALFA-mNG-hRft1-3xFLAG-Ist2^590-946^ to the cER (Fig. 4D, top). Analysis of the fluorescence distribution (fraction of fluorescence in nER versus total fluorescence per cell) revealed that the redistributions were quantitatively significant (Fig. 4E). These data indicate that hRft1 can be redistributed within the ER by appending cytoplasmic tethering modules to the N- and C-termini, indicating that these termini are oriented to the cytoplasm.

Both the AlphaFold2 model and an HHpred search (56) of the Protein Data Bank indicate that hRp1 has a fold resembling that of members of the multidrug/oligosaccharidyl- lipid/polysaccharide (MOP) family of transporters (57), with structural homology to bacterial MurJ lipid II flippases which are proposed to operate by an alternating access mechanism powered by membrane potential (58–61). The Alphafold model shows hRp1 in an inward-open conformation (Fig. 4F). The N- and C-terminal lobes define a central hydrophilic region corresponding to a putative substrate binding pocket which is 2.3 nm wide at the membrane-water interface and open to the cytoplasm (Fig. 4F). We used the ConSurf bioinformatics tool (62, 63) to investigate the evolutionary conservation of residues in hRp1 and found high conservation in this hydrophilic region (Fig. 4G). Most of the known RFT1-CDG disease mutations (Table 3) mapped to this region, an interesting exception being the R37 residue (disease variant R37L, Table 3) which is located on the luminal side of the protein where it is potentially involved in stabilizing the interaction between the N- and C-terminal lobes in the modelled conformation (Fig. 4F).

Several charged residues (R37, R63, R290, R442, E260) are fully conserved in Rft1 sequences (Fig. 4G, ConSurf scores of 8 or greater), and these are mostly located in the hydrophilic central region. Arginine residues are also found in the central cavity of MurJ where they are required for function (59). The TM1 helix is strongly amphipathic with a conserved Q4-R8-F12-N15 motif, the three hydrophilic residues (Q21, R25, N32) clustering to one side of the helix (57). Using evolutionary coupling analysis (58, 64, 65), we identified A155 as a potentially interesting residue. Although this residue is not highly conserved (ConSurf grade = 6), the PROVEAN protein webserver (66) indicates that a charge substitution, A155E or A155K, would be deleterious to function.

### Functional test of hRft1 variants with point mutations

We next chose to develop a structure-function model of hRft1, using the yeast system to test the functional consequences of introducing mutations at key sites. We chose a number of the sites described above (Fig. S3A) for mutagenesis; we also included several RFT1-CDG mutations for analysis (including R67C, which was the first RFT1-CDG mutation to be identified (15)). Using a plasmid shuffling approach we tested the ability of the hRft1 point mutants to support growth. Thus, we transformed KSY512 cells with *HIS3* plasmids encoding hRft1-3xFLAG point mutants under control of the GPD promoter, then tested the cells for growth on 5-FOA plates. Figure S3B shows that of the 16 mutants that we tested, R25W, G276D, R290A, Y301C and R442A failed to support growth. To determine whether this was a result of low protein expression, we transformed wild-type cells with the same plasmids and analyzed the expression level of the constructs by immunoblotting. Figure S3C (quantification in Fig. S3D) shows that these variants are expressed at reasonable levels compared with most of the other constructs, indicating that their phenotype is likely directly due to loss of function. Surprisingly, the Q21L and R37A mutants were poorly expressed yet supported growth (Fig. S3).

To obtain more quantitative information, we made use of the KSY512 cells that we recovered from the plasmid shuffling experiment (Fig. S3B); these cells express hRft1 variants that retain sufficient functionality to enable growth. Quantitative measurements of cell growth over a 36-h period (exemplary growth curves are provided in Figure 5A) showed that growth was slowed significantly in several strains (Fig. 5C). Immunoblotting analysis (Fig. 5B, bottom panel (anti-FLAG blot) and Fig. 5D) indicated that all proteins were reasonably expressed in comparison with wild-type protein, except for R37 which was expressed at a very low level as noted above (Fig. S3C, D). We next assessed the steady state glycosylation status of CPY in these strains (Fig. 5B, E). Compared with cells expressing the wild-type protein which had a CPY Glycoscore of ∼70, CPY was hypoglycosylated in all strains expressing the hRft1 mutants, with Glycoscores in the range 50-60 (Fig. 5E), with the exception of G340S which had a CPY Glycoscore similar to that in wild-type cells. Closer examination of these data (Fig. 5F-H) revealed several points of interest. Faster growing strains had generally higher CPY Glycoscores, except for the strain expressing E260A which grew slowly despite reasonable glycosylation (Fig. 5F). The R37A variant appeared to be hyperactive (Fig. 5G, H): it was poorly expressed (Fig. S3C, D and Fig. 5B, D) yet enabled normal cell growth (Fig. 5A, C) and supported an average level of CPY glycosylation (Glycoscore ∼60) (Fig. 5E). Because R37 may stabilize the interaction between the N- and C-terminal lobes in the inward-open conformation of hRft1 (see above)(Fig. 4F), a mutation at this site may result in reduced stability, favoring conformational switching. R63A had the poorest CPY Glycoscore of the mutants that we tested, despite a normal expression level, and the disease mutant R67C seemed to be functionally compromised — it was the most highly expressed of all the variants we tested, including the wild-type protein, yet cells expressing this variant underperformed in terms of doubling time and CPY Glycoscore (Fig. 5G, H). Our results on the R67C mutant differ from those presented in a previous report (15) where a colony sectoring assay was used to compare wild-type hRft1 and hRft1(R67C) in yeast. This assay indicated that the R67C mutant was non-functional as no colony sectoring could be observed.

**Figure 5.**
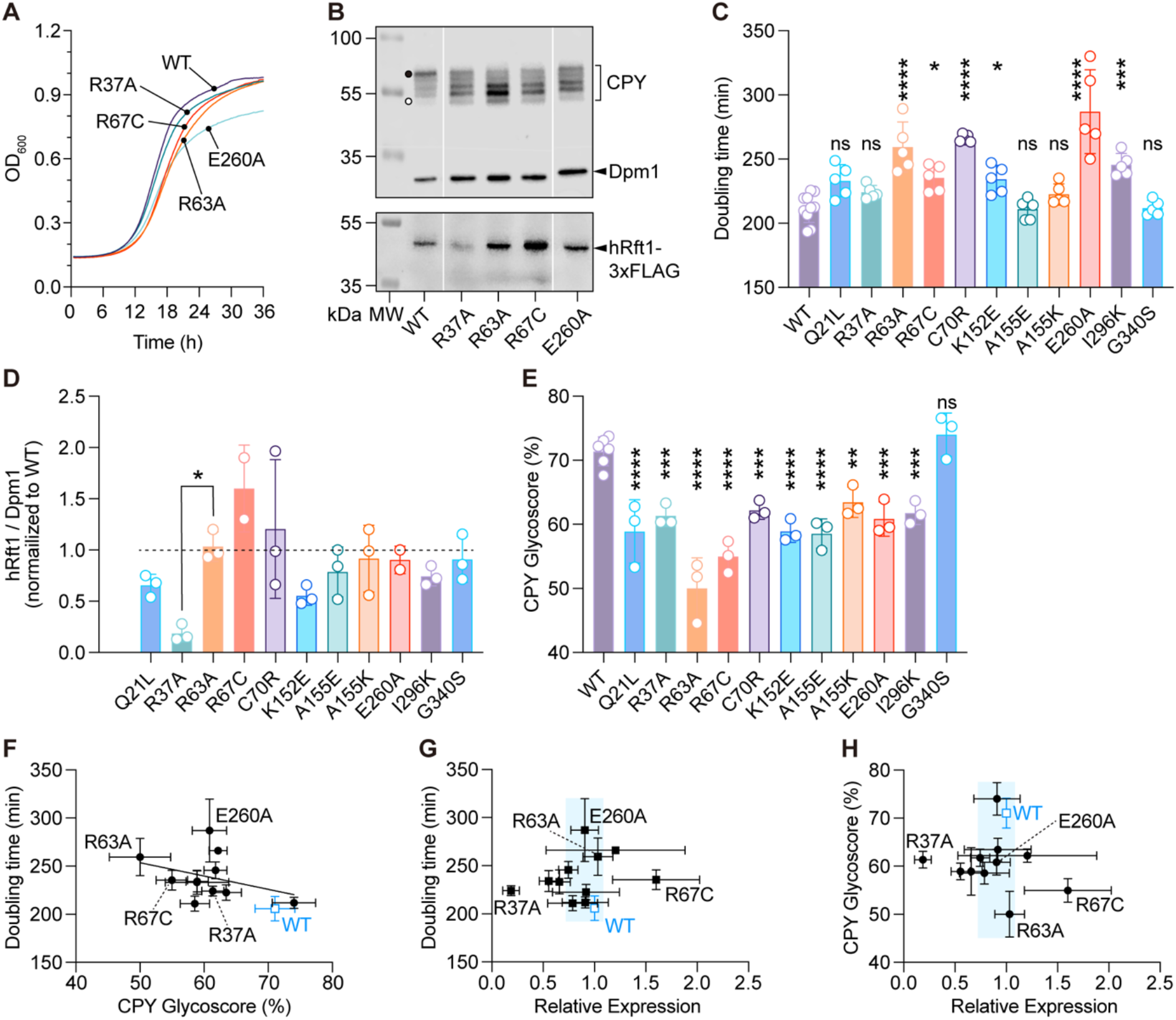
Analysis of cells expressing hRft1 point mutants. Plasmid shuffling was used to replace the hRft1-expressing *URA3* plasmid in KSY512 cells with *HIS3* plasmids expressing wild-type hRft1-3xFLAG or corresponding point mutants. A. KSY512 cells expressing hRft1-3xFLAG variants were cultured in SD(-His) medium to mid-log phase and diluted to OD_600_=0.01. OD_600_ was measured every 15 min for 36 h in a plate reader. The measurement was repeated 5 times and average data are presented. B. KSY512 cells expressing hRft1 point mutants were cultured in SD (-His) medium to log-phase, harvested, and analyzed by SDS-PAGE and immunoblotting with anti-CPY, anti-Dpm1, and anti-FLAG antibodies. The black dot and white dot in the CPY blot indicate fully glycosylated and non-glycosylated CPY, respectively. C. Doubling time was determined from the exponential phase of growth curves, including those shown in panel A. Data are shown as a bar chart (mean ± S.D. (n=5 technical replicates)) with individual values (*p=0.0229 (R67C) and 0.0359 (K152E), ***p=0.0003, ****p<0.0001, ns, not significant, ordinary one-way ANOVA with all samples compared with WT). D. hRft1-3xFLAG protein expression levels were quantified by calculating the ratio of the intensity of the hRft1-3xFLAG band to Dpm1 and normalizing to that of the WT sample from immunoblots such as the one shown in panel B. Data are presented as mean ± S.D. (n=3 biological replicates), with individual values indicated. Ordinary one-way ANOVA revealed no significant differences between the expression level of the mutants in comparison with R63A (chosen as reference because its expression was similar to that of wild-type hRft1-3xFLAG), except for R37A indicated as *, p=0.0138. E. The intensity of each CPY band in panel B was analyzed and quantified to obtain the CPY Glycoscore. Data are presented as mean ± S.D. (n=3 biological replicates)(**p=0.0044, ***p=0.0003 (R37A), 0.0008 (C70R), 0.0001 (E260A), 0.0005 (I296K),****p<0.0001, ns, not significant, ordinary one-way ANOVA with all samples compared with WT). F. Correlation between doubling time and CPY Glycoscore. Data points are mean ± S.D. (n=5 for doubling time, n=3 for CPY Glcosycore). G. Correlation between doubling time and expression. Data points are mean ± S.D. (n=5 for doubling time, n=3 for expression). H. Correlation between CPY Glycoscore and expression. Data points are mean ± S.D. (n=3). G-H. The light blue shaded box is centered on the median of all expression values (x-axis) with a width +/- 20% of the median. The height of the box covers the y-axis data range.

To determine if the mutant proteins were correctly localized to the ER, we integrated N-terminal mNG-tagged variants of R37A, R63A, R67C and E260A into yeast cells. Fluorescence microscopy revealed a typical ER pa{ern for all constructs, except for the presence of occasional varicosities in the cortical ER, seen in over half the R63A and R67C-expressing cells (Fig. S4).

### Concluding remarks

Rft1 is nearly ubiquitously found in eukaryotes (67) where it plays a critical yet undefined role in protein *N*-glycosylation. Despite its importance, a basic molecular characterization of the protein is not currently available, and this is what we have provided in this paper. We show that hRft1 is a polytopic ER membrane protein (Figs. 1E, 4A), with its N- and C-termini oriented to the cytoplasm (Fig. 4C-E). It is not *N*-glycosylated (Fig. 4B). The AlphaFold2 model of hRp1 (Fig. 4F) resembles that of the inward-open conformation of an alternating access transporter (68), with a hydrophilic cavity open to the cytoplasm (Fig. 4F). The cavity, which likely represents a substrate binding site, contains several charged residues that are conserved in all Rp1 sequences and which we demonstrate to be functionally important by quantifying the ability of the corresponding point mutants to support cell growth and *N*-glycosylation in the yeast model system (Figs. 5, S3B).

Rft1 belongs to the multidrug/oligosaccharidyl-lipid/polysaccharide (MOP) exporter superfamily of transporters (57) which includes the bacterial MurJ flippases (59). MurJ proteins export Lipid II, a cytoplasmically synthesized undecaprenyl diphosphate-linked peptidoglycan building block, across the membrane to the periplasmic side (3, 59, 69). They are proposed to operate by an alternating access mechanism (68) in which transbilayer export of Lipid II is coupled to the movement of a counterion down its electrochemical gradient (61). The structural similarity between Rp1 and MurJ, and between M5-DLO and Lipid II (3, 59), revives the possibility that Rft1 may play a role in translocating M5-DLO across the ER membrane for *N*-glycosylation. However, many points need to be considered in order to develop this hypothesis.

First, in contrast to MurJ-catalyzed lipid export which requires membrane potential (61), transbilayer translocation of M5-DLO and other isoprenoid-based lipids in the ER appears to be mediated by scramblases which equilibrate lipids between the two leaflets of the bilayer independent of metabolic energy inputs (5, 51, 70). It is possible that Rft1 acts as a passive or equilibrative transporter, moving M5-DLO down the transbilayer concentration gradient generated via its synthesis on the cytoplasmic side of the ER and consumption on the lumenal side (Fig. 1A). It is also possible that Rft1 may operate by a mechanism that is unrelated to the canonical alternating access model. Of relevance to this idea, a recent report (71) showed that the lactose permease, LacY, a well-characterized alternating access symporter (72), scrambles phospholipids independently of its substrates (proton, lactose) which drive the conformational changes associated with alternating access.

Second, as noted in the Introduction and shown in Fig. 3D, Rft1 is not necessary for M5-DLO scrambling in cell-free systems where specific scrambling is primarily accomplished by another protein or protein complex in the ER (5, 21–23). Consistent with this point, it was previously reported that M5-DLO scramblase activity is associated with ER glycoproteins or glycoprotein complexes (51) which rules out a role for Rft1 which we show here is not glycosylated (Fig. 4B). Whereas these results do not rule out a transport function for Rft1 — something that can be tested in the future by assaying purified protein in a reconstituted system — they leave open the question of the precise nature of the lipid or other substrate being transported and whether this transport function underlies Rft1’s essentiality in yeast and mammalian cells. A detailed investigation of these points awaits future work.

## EXPERIMENTAL PROCEDURES

### Plasmids

The plasmids used in this paper are listed in Table 1. Plasmids were propagated using DH5-α *E. coli* cells which were cultured at 37°C and 210 rpm in LB+Amp medium (1% tryptone, 0.5% yeast extract, 1% NaCl, and 100 µg/ml ampicillin). Plasmid construction was as follows:

EcAKM135 (hRft1-3xFLAG), EcAKM141 (hRft1^R67C^-3xFLAG) and EcAKM187 (p416-P_GPD_-hRFT1-3xFLAG): hRft1-3xFLAG and hRft1^R67C^-3xFLAG fragments were PCR amplified from plasmids AP63 and AP65, respectively, and inserted into BamHI-HindIII-digested p413*-P_GPD_* or p416*-P_GPD_* vector using the NEBuilder HiFi DNA Assembly kit (New England Biolabs, E5520S).

EcAKM147 (P_GPD_-ALFA-mNG-hRft1-3xFLAG): the mNeonGreen fragment was amplified from pFA6a-2xmNeonGreen-kanMX and inserted into EcAKM135 using the NEBuilder HiFi DNA Assembly kit to construct EcAKM136 (mNG-hRft1-3xFLAG). Next, the ALFA-tag was inserted between *P_GPD_* and mNeonGreen by PCR-based site-directed mutagenesis.

EcAKM148 (P_TEF_-ALFA-mNG-hRft1-3xFLAG): the *P_GPD_* promotor of EcAKM136 was replaced with the *P_TEF_* promotor from p413-*P_TEF_* by traditional cloning methods using XbaI and XhoI to generate the necessary fragments. The ALFA-tag was inserted as above.

EcAKM149 (P_ADH_-ALFA-mNG-hRft1-3xFLAG): the *P_GPD_* promotor of EcAKM147 was replaced with the *P_ADH_* fragment digested from p415*-P_ADH_* with SacI and XbaI.

EcAKM150-152 (pCfB2195-P_GPD/TEF/ADH_-ALFA-mNG-hRft1-3xFLAG): *P_GPD/TEF/ADH_*-ALFA-mNeonGreen-hRft1-3xFLAG fragments were PCR amplified from EcAKM147-149 and cloned into pCfB2195 digested with AsiSI using the NEBuilder HiFi DNA Assembly kit.

EcAKM153 (pRG205MX-mCherry-HDEL) and EcAKM154 (pRG205MX-P_ADH_-Nvj1^1-121^-ALFA-nB): mCherry-HDEL and *P_ADH_-*Nvj1^1-121^-ALFA-nB were inserted into pRG205MX. Insertion and vector fragments were created by digesting pJF132 (pRS305-mCherry-HDEL), FFP386 (pRS405-P_ADH_-Nvj1^1-121^-ALFA-nB), and pRG205MX, respectively, with SacI and XhoI. The AscI site in the ADH terminator region of EcAKM153 was mutated by PCR-based site-directed mutagenesis.

EcAKM216 (pCfB2195-P_GPD_-mNG-Alg2) and EcAKM217 (pCfB2195-P_GPD_-mNG-Alg14): the hRft1-3xFLAG sequence of EcAKM170 (p416-P_GPD_-mNG-hRft1-3xFLAG) was replaced with Alg2 or Alg14 fragments amplified from EcAKM197 (pGB1805-Alg2) and EcAKM198 (pGB1805-Alg14), respectively. Then *P_GPD_*-mNG-Alg2 or *P_GPD_*-mNG-Alg14 fragments were PCR amplified and cloned into pCfB2195 digested with AsiSI using the NEBuilder HiFi DNA Assembly kit.

*RFT1* mutants were created by PCR-based site-directed mutagenesis using EcAKM135 (hRft1- 3xFLAG) or EcAKM150 (pCfB2195-P_GPD_-ALFA-mNG-hRft1-3xFLAG) as templates.

EcAKM222 (pRS315-yRft1): yeast *RFT1* gene fragment including 500bp upstream and 400bp downstream of its open reading frame was PCR amplified from yeast genome and cloned into pRS315 vector digested with SmaI using the NEBuilder HiFi DNA Assembly kit. To construct EcAKM221 (yRft1-3xFLAG), yRft1 fragments were PCR amplified from EcAKM222 and inserted into BamHI-HindIII-digested p413*-P_GPD_* vector using the NEBuilder HiFi DNA Assembly kit.

EcAKM218 (pCfB2195-P_GPD_-ALFA-mNG-yRft1-3xFLAG) and EcAKM219 (pCfB2195-P_ADH_-ALFA-mNG-yRft1-3xFLAG): hRft1 fragment in EcAKM150 was replaced with yRft1 from EcAKM222 by PCR cloning method using the NEBuilder HiFi DNA Assembly kit. *P_GPD_-*ALFA-mNG-yRft1-3xFLAG fragment was PCR amplified from EcAKM218 and cloned into PCR amplified p413*-P_GPD_* vector to construct EcAKM220 (P_GPD_-ALFA-mNG-yRft1-3xFLAG).

EcAKM224 (pCfB2195-P_ADH_-ALFA-mNG-hRft1-3xFLAG-Ist2^590-946^): the C-terminal sequence of *IST2* (590-946) was PCR amplified and cloned into PCR amplified EcAKM152 (pCfB2195-P_ADH_-ALFA- mNG-hRft1-3xFLAG) vector using the NEBuilder HiFi DNA Assembly kit.

### Yeast strains and culture conditions

The strains used in this paper are listed in Table 2. Yeast cells were cultured in YPD (1% yeast extract, 2% peptone, and 2% dextrose), synthetic defined (SD) or synthetic complete (SC) (0.17% yeast nitrogen base without amino acids and ammonium sulfate, 0.5% ammonium sulfate, and 2% dextrose with appropriate amino acids and bases as necessary). Cells were cultured at 30°C and 250 rpm. 5-Fluoroorotic acid (5-FOA; ZYMO RESEARCH F9003), or doxycycline (Sigma D9891), were included as indicated. YPO (1% yeast extract, 2% peptone, 0.1% dextrose, 0.125% oleic acid, 0.5% Tween80) medium was used to induce lipid droplet formation.

**Table 2.** Yeast strains.

| YEAST STRAIN NAME | SYSTEMATIC NAME | GENOTYPE | SOURCE / REFERENCE |
| --- | --- | --- | --- |
| Wild-type | BY4741 | *MATa his3Δ1 leu2Δ0 met15Δ0 ura3Δ0* | (1) |
| Wild-type | S288C | *MATα SUC2 gal2 mal2 mel flo1 flo8-1 hap1 ho bio1 bio6* | Gift from Maya Schuldiner (Weizmann Institute of Science)(2) |
| *rft1::KANMX4 / RFT1* | YAKM158 | BY4743, *his3Δ1 leu2Δ0 ura3Δ0 rft1::KANMX4 / RFT1* | Yeast Knockout (YKO) Collection (3) |
| KSY512 | KSY512 / YAKM188 | BY4741, *rft1::KANMX4* carrying p416-P<sub>GPD</sub>-hRft1-3xFLAG (EcAKM187) | This study |
| mCherry-HDEL | YAKM223 | S288C, *mCherry-HDEL-LEU2MX* | This study |
| P<sub>GPD</sub>-mNG-hRft1<br>mCherry-HDEL | YAKM225 | YAKM223, *P<sub>GPD</sub>-ALFA-mNeonGreen-hRFT1-3xFLAG-hphMX* | This study |
| P<sub>TEF</sub>-mNG-hRft1<br>mCherry-HDEL | YAKM224 | YAKM223, *P<sub>TEF</sub>-ALFA-mNeonGreen-hRFT1-3xFLAG-hphMX* | This study |
| P<sub>ADH</sub>-mNG-hRft1<br>mCherry-HDEL | YAKM234 | YAKM223, *P<sub>ADH</sub>-ALFA-mNeonGreen-hRFT1-3xFLAG-hphMX* | This study |
| Erg6-mCherry | ATY502/YAKM187 | BY4741, *ERG6-mCherry::HIS* | Gift from Alexandre Toulmay (University of Texas Southwestern Medical Center) |
| P<sub>tet</sub>-Rft1 | YAKM147 | *URA::CMV-tTA, his3-1 leu2-0 met15-0 RFT1::kanR-TetO<sub>7</sub>-CYC1<sub>TATA</sub>-RFT1* | Gift from Maya Schuldiner (Weizmann Institute of Science)(4) |
| P<sub>tet</sub>-Rft1 P<sub>ADH</sub>-AFLA-mNG-hRft1 | YAKM172 | YAKM147, *P<sub>ADH</sub>-ALFA-mNeonGreen-hRFT1-3xFLAG-hphMX* | This study |
| P<sub>tet</sub>-Rft1 Nvj1-nB P<sub>ADH</sub>-AFLA-mNG-hRft1 | YAKM173 | YAKM172, *P<sub>ADH</sub>-NVJ1<sup>1-121</sup>-ALFA-nB-LEU2MX* | This study |
| P<sub>ADH</sub>-mNG-hRft1<br>Erg6-mCherry | YAKM235 | ATY502, *P<sub>ADH</sub>-ALFA-mNeonGreen-hRFT1-3xFLAG-hphMX* | This study |
| P<sub>GPD</sub>-mNG-hRft1<sup>R37A</sup><br>mCherry-HDEL | YAKM248 | YAKM223, *P<sub>GPD</sub>-ALFA-mNeonGreen-hRFT1<sup>R37A</sup>-3xFLAG-hphMX* | This study |
| P<sub>GPD</sub>-mNG-hRft1<sup>R63A</sup><br>mCherry-HDEL | YAKM249 | YAKM223, *P<sub>GPD</sub>-ALFA-mNeonGreen-hRFT1<sup>R63A</sup>-3xFLAG-hphMX* | This study |
| P<sub>GPD</sub>-mNG-hRft1<sup>R67C</sup><br>mCherry-HDEL | YAKM250 | YAKM223, *P<sub>GPD</sub>-ALFA-mNeonGreen-hRFT1<sup>R67C</sup>-3xFLAG-hphMX* | This study |
| P<sub>GPD</sub>-mNG-hRft1<sup>E260A</sup><br>mCherry-HDEL | YAKM251 | YAKM223, P<sub>GPD</sub>-ALFA-mNeonGreen-hRFT1<sup>E260A</sup>-3xFLAG-hphMX | This study |
| P<sub>GPD</sub>-mNG-yRft1<br>mCherry-HDEL | YAKM301 | YAKM223, P<sub>GPD</sub>-ALFA-mNeonGreen-yRFT1-3xFLAG-hphMX | This study |
| P<sub>ADH</sub>-mNG-yRft1<br>Erg6-mCherry | YAKM302 | ATY502, P<sub>ADH</sub>-ALFA-mNeonGreen-yRFT1-3xFLAG-hphMX | This study |
| P<sub>GPD</sub>-mNG-Alg2<br>Erg6-mCherry | YAKM274 | ATY502, P<sub>GPD</sub>-ALFA-mNeonGreen-ALG2-3xFLAG-hphMX | This study |
| P<sub>GPD</sub>-mNG-Alg14<br>Erg6-mCherry | YAKM275 | ATY502, P<sub>GPD</sub>-ALFA-mNeonGreen-ALG14-3xFLAG-hphMX | This study |
| P<sub>tet</sub>-Rft1 P<sub>ADH</sub>-AFLA-mNG-hRft1-Ist2<sup>590-946</sup> | YAKM287 | YAKM147, P<sub>ADH</sub>-ALFA-mNeonGreen-hRFT1-3xFLAG-IST2<sup>590-946</sup>-hphMX | This study |

**Table 3.** Known muta0ons in R41 associated with RFT1-CDG.

| MUTATION | REFERENCE |
| --- | --- |
| R25W | (1) |
| R37L/R442Q | (2) |
| I43V | (3) |
| R67C | (4, 5) |
| C70R | (1) |
| K152E | (2, 5-7) |
| G276D | (1) |
| I296K/I296R | (6) |
| E298K | (5) |
| Y301C | (8) |
| G340S | (9) |
| R442Q/M408V | (10) |

Strain KSY512 was generated from a heterozygous *rft1::KANMX4* / *RFT1* deficient strain (YAKM158) which was transformed with EcAKM187 (p416-P_GPD_-hRft1-3xFLAG) and sporulated. Individual spores were obtained by tetrad analysis and colonies, initially grown on YPD, were tested for growth on YPD+G418 and SD(-Ura) plates. The disruption of genomic *RFT1* in KSY512 was also checked by PCR with *KANMX4*-specific primers.

YAKM223 (mCherry-HDEL) and YAKM173 (P_tet_-Rft1 Nvj1-nB P_ADH_-ALFA-mNG-hRft1) were created by transforming S288C or YAKM172 yeast strains with EcAKM153 and EcAKM154 plasmids, respectively, digested with AscI. After transformation, colonies were selected by SD(-Leu) plate.

To integrate mNG-Alg2, mNG-Alg14, ALFA-mNG-yRft1-3xFLAG, ALFA-mNG-hRft1-3xFLAG, ALFA- mNG-hRft1-3xFLAG-Ist2^590-946^, and point mutants under control of GPD, TEF or ADH promoters into different yeast strains (YAKM223, ATY502, YAKM147), the cells were transformed with plasmids EcAKM150-152,164-167, 216-219, 224 that were linearized with NotI. After integration, colonies were selected on YPD containing 300 µg/ml hygromycin B (Gibco 10-687-010).

### Yeast growth assay

For plate assays, cells were adjusted to OD_600_=1.0 and 10-fold serial dilutions were spotted. Plates were incubated at 30°C for 2-3 days before being photographed. For generating growth curves, cells were cultured in SD(-His) medium to mid-log phase and diluted to OD_600_=0.01 in a 96-well plate. The plate was covered with a Breathe-Easy® polyurethane sealing membrane (Sigma Z380059) and placed in a SpectraMax i3x plate reader (Molecular Devices), incubated at 30°C, and OD_600_ was measured every 15 min for 36 h. The plate was shaken for 5 sec before each measurement. Doubling time was calculated as (t_2_-t_1_) x log2/(logC_2_-logC_1_), where the OD_600_ of the culture at times t_1_ and t_2_ correspond to C_1_ and C_2_, respectively. The values of C_1_ and C_2_ were chosen to be ≥0.35 and ≤0.5, respectively.

### SDS-PAGE immunoblotting

Cells were cultured to mid-log phase, and 2 OD_600_ units were collected. The cells were lysed in 0.28 ml 0.2 M NaOH, 0.5% β-mercaptoethanol and incubated on ice for 5 min before adding 1 ml 15% TCA to precipitate all proteins. After 10 min incubation on ice, precipitated proteins were collected by centrifugation (14,000 rpm, 5 min, 4°C), washed with 0.6 ml ice-cold acetone, air-dried, and solubilized overnight in 0.2 ml 2% SDS, 5 mM NaOH, 2% β-mercaptoethanol. The samples were mixed with 100 μl of 2xSDS-PAGE loading buffer and used for SDS-PAGE. CPY, Dpm1, and FLAG were detected by immunoblotting with primary anti-CPY (1:5000; Invitrogen 10A5B5), anti-Dpm1 (1:1000; Life technology 5C5A7)), or anti-FLAG (1:2000; Sigma F1804) antibodies, followed by HRP-conjugated secondary anti-mouse antibody (1:5000, Promega W402B), diluted as indicated. Images were captured on an Odyssey XF (LI-COR) and band intensities were calculated using Image Studio software (LI-COR).

CPY glycosylation score calculation was described previously (29). Briefly, the intensities of each CPY band were quantified using Image Studio software, and the data set was used to calculate the CPY Glycoscore. The relative intensities of bands were multiplied by 4, 3, 2, 1, and 0, representing fully glycosylated CPY, and lacking 1, 2, 3, and 4 *N*-glycans, respectively. Values were summed, divided by 400, and converted to percentages.

### Fluorescence microscopy

Cells were cultured in YPD medium to mid-log phase, harvested, washed with water, and collected by centrifugation at room temperature in a microcentrifuge. To induce lipid droplet formation, mid-log phase cells were harvested and washed with YPO medium. Cells were resuspended with YPO medium and incubated at 30°C for 16 hours, then washed with water and collected by centrifugation at room temperature in a microcentrifuge. Cells were imaged using an LSM 880 Confocal Laser Scanning Microscope (Zeiss) with ZEN Microscopy software (Zeiss), 63x lens (Plan-Apochromat 63x/1.4 Oil (Zeiss)), and 488 nm and 561 nm lasers for mNeonGreen and mCherry, respectively. Alternatively, mNeonGreen images were taken with a Nikon, Eclipse Ti2 microscope with a 60x lens (Plan Apo λ 60x/1.40 Oil (Nikon)) using NIS-Elements (Nikon) software and a GFP filter. For grayscale images, acquired images were processed with ImageJ software by selecting “Images > Lookup tables > Grays” and “Edit > Invert”. For data presented in Figure 4D, E, fluorescence was quantified using Image J as follows: the total fluorescence (F_total_) and nuclear fluorescence (F_nuc_) of each cell was determined within approximately circular outlines of the cell and the nucleus. Similar measurements were made using the same outlines placed in a cell-free area of the image to determine background correction.

### Preparation of yeast microsomes and the Triton Extract (TE)

Salt-washed microsomes were prepared from a homogenate of KSY512 yeast cells as described previously (22), except an additional centrifugation step (20,000 × *g_av_*, 30 min, 4°C) was included after the initial low-speed spin to clear the homogenate of mitochondria, assessed by immunoblotting to detect Por1, the mitochondrial outer membrane porin. The salt-washed microsomes were incubated with ice-cold Triton X-100 as previously described (22) to generate a Triton extract (TE, buffer composition: 50 mM Hepes, pH 7.4, 150 mM NaCl, 1% (w/v) Triton X-100) selectively enriched in ER membrane proteins (5). hRft1-3xFLAG was quantitatively eliminated from the TE by incubating the extract (300 μl) with anti-FLAG M2 affinity gel resin (Sigma A220; bed volume 100 μl); an equivalent aliquot of TE was mock-treated in parallel. Specific elimination of hRft1-3xFLAG was confirmed by immunoblotting with anti-FLAG and anti-Dpm1 antibodies as described above. The protein concentration of the TE, irrespective of treatment, was ∼0.5 mg/ml as determined by the Pierce Micro BCA Protein Assay kit (ThermoFisher 23235).

### Reconstitution of proteoliposomes and scramblase assays

Proteoliposomes and protein-free liposomes were reconstituted using a previously described one-pot protocol (22) with some modifications. Briefly, to prepare three reconstituted samples, 10 mg egg phosphatidylcholine (Avanti Polar Lipids 840051), 45 μg 16:0-06:0 NBD-PC (Avanti Polar Lipids 810130C) and ∼40,000 cpm [^3^H]M5-DLO (22) were added from chloroform or chloroform/methanol stock solutions to a glass tube, dried under a stream of nitrogen, dissolved in 1 ml pentane (Sigma 34956-1L) and dried again under nitrogen. The resulting lipid film was dissolved to clarity by stepwise addition of 10/100/1 buffer (10 mM Hepes, pH 7.4, 100 mM NaCl, 1% (w/v) Triton X-100) to a total volume of 1.85 ml. The lipid solution was distributed into three 2-ml Eppendorf tubes (600 μl per tube) and supplemented with 400 μl of either 10/100/1 buffer, mock-treated TE, or α-FLAG-treated TE to generate protein-free liposomes, complete proteoliposomes, or Rft1-deficient proteoliposomes, respectively. The samples were incubated at room temperature with end-over-end mixing for 30 min before adding washed BioBeads SM2 (Bio-Rad 152-39120) in two stages. First, each sample received 100 mg BioBeads and was incubated with end-over-end mixing at room temperature for 3 h. Next, the samples were supplemented with 200 mg BioBeads and incubated with end-over-end mixing in a cold-room overnight (∼15 h). The samples were removed from the BioBeads and centrifuged in a Beckman TLA100.2 rotor at 75,000 rpm (∼250,000 x *g_max_*)at 4°C for 1 h. The resulting membrane pellets were each resuspended in 200 μl MMC buffer (10 mM Hepes, pH 7.4, 100 mM NaCl, 3 mM MgCl_2_,1 mM MnCl_2_,1 mM CaCl_2_). Aliquots of the resuspended pellets were taken for dynamic light scattering and phospholipid scramblase activity assays as described (46), and M5-DLO scramblase assay according to a standard protocol (22).

## Data availability

Plasmid sequences are deposited on Zenodo at https://zenodo.org/records/12100636. All other data are contained within the manuscript.

## Supporting information

This article contains supporting information (Figs. S1-S4 and Tables 1-3).

## Acknowledgements

We thank the Optical Microscopy and Imaging services of the Microscopy & Image Analysis Core Facility at Weill Cornell, as well several individuals at Weill Cornell for use of their equipment: James Jordan and Baran Ersoy (plate reader), Beate Schwer (tetrad dissecting microscope), and David Simon (fluorescence microscope). We thank Ralf Erdmann, Roger Schneiter and Vineet Choudhary for advice on induction of lipid droplets in yeast and the following individuals for reagents: Alice Verchère for [^3^H]M5-DLO, Maya Schuldiner (Weizmann Institute) for Tet-off strains, Alexandre Toulmay and Jonathan Friedman (University of Texas Southwestern Medical Center) for the Erg6-mCherry-expressing cells and the mCherry-HDEL plasmid, respectively, Bianca Esch and Florian Fröhlich (Osnabrück University) for reagents to carry out nanobody-based recruitment of Rft1 to ER domains, Liesbeth Veenhoff ((European Research Institute for the Biology of Aging, University of Groningen) for an Ist2 plasmid, and Hiroki Okada (Erfei Bi laboratory, University of Pennsylvania) for integration plasmids.

## Funding and additional information

We acknowledge support from National Institutes of Health grants R21 HD109719 and R01 GM146011 (A.K.M.), and the Fonds National de la Recherche Scientifique (FRS-FNRS; Grant number 1.B.089.24F)(G.N.C.). The content is solely the responsibility of the authors and does not necessarily represent the official views of the National Institutes of Health.

## Footnotes

The abbreviations used are: 5-FOA, 5-fluoroorotic acid; ADH, constitutively active promoter from alcohol dehydrogenase; CDG, congenital disorder of glycosylation; CPY, carboxypeptidase Y; DDM, n-dodecyl-β-maltoside; DLO, dolichol-linked oligosaccharide; DLS, dynamic light scattering; Dox, doxycyline; ECL, extracellular loop; ER, endoplasmic reticulum; G3M9-DLO, Glucose_3_Mannose_9_GlcNAc_2_-diphosphate-dolichol; Glc, glucose; GlcNAc, *N*-acetylglucosamine; Glycoscore, glycosylation score; GPD, strong, constitutively active promoter from glyceraldehyde 3-phosphate dehydrogenase; His, histidine; hRft1, human Rft1; ICL3, intracellular loop 3; LD, lipid droplet; LUV, large unilamellar vesicle; M5-DLO, Mannose_5_GlcNAc_2_-diphosphate-dolichol; Man, mannose; mCPY, mature CPY; mNG, mNeonGreen; nB, nanobody; NBD, 7-nitro-2,1,3- benzoxadiazole; OST, oligosaccharyltransferase; PC, phosphatidylcholine; SD medium, synthetic defined medium; TE, Triton X-100 Extract enriched in ER membrane proteins; Ura, uracil

---

**Supplementary Figure S1.**
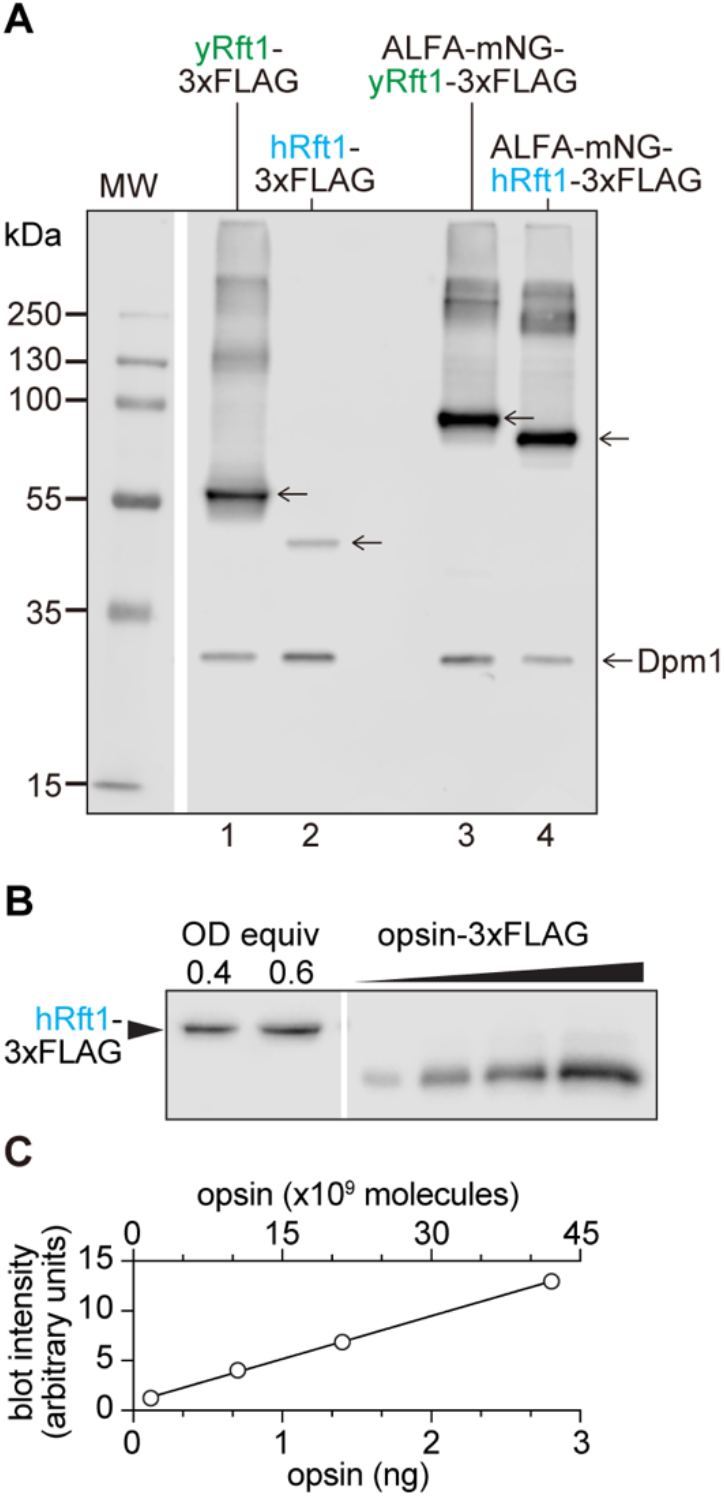
Quantifying expression of hRft1 constructs. A. Expression of yRft1 and hRft1 proteins. Plasmid shuffling was used to replace hRft1-3xFLAG in KSY512 cells with yRft1-3xFLAG, yRft1-3xFLAG, ALFA-mNG-yRft1-3xFLAG, and ALFA-mNG-hRft1-3xFLAG expressed under control of the GPD promoter. Cells were cultured to log-phase, harvested, and analyzed by SDS-PAGE and immunoblotting with anti-FLAG and anti-Dpm1 (loading control) antibodies. Unmarked arrows indicate Rft1-3xFLAG and ALFA-mNG-mNG-Rft1-3xFLAG. All Rft1 constructs migrate faster than expected based on their predicted molecular weights; yRft1 is approximately 6 kDa larger than hRft1. The relative expression levels of yRft1 and hRft1 constructs is as follows (mean ± S.D., n=3 biological replicates): yRft1-3xFLAG/ yRft1-3xFLAG = 6.7 ± 2.4 and ALFA-mNG-yRft1-3xFLAG/ ALFA-mNG-hRft1-3xFLAG = 1.2 ± 0.5. B. KSY512 cells expressing hRft1-3xFLAG were cultured to log-phase and proteins (corresponding to 0.4 and 0.6 OD equivalents of cells) were analyzed by SDS-PAGE alongside 3xFLAG-tagged bovine opsin standards (0.12, 0.70, 1.4, and 2.8 ng (obtained by diluting a 117 ng/µL stock solution)(Ploier et al. (2016) Nat Commun 7, 12832)). FLAG-tagged hRft1 and opsin were visualized by immunoblotting with anti-FLAG antibodies. The opsin band runs just above the 35 kDa molecular weight marker; hRft1 runs between the 55 and 35 kDa markers as seen in panel A, lane 2. C. The immunoblot signal intensity for the opsin standards (obtained by densitometry using ImageJ) was graphed against the protein amount loaded onto the gel (bottom x-axis), and linear regression was used to obtain a calibration plot. The x-axis at the top of the plot was obtained by converting opsin mass to number of molecules using a molecular weight of 40,000 g/mole. This plot was used to convert the Rft1 signal intensity (3.8 and 5.1 units (obtained using ImageJ), corresponding to 0.4 and 0.6 OD equivalents, respectively) to number of molecules, yielding a final result of 840 ± 35 molecules/cell (n=2 technical replicates).

**Supplementary Figure S2.**
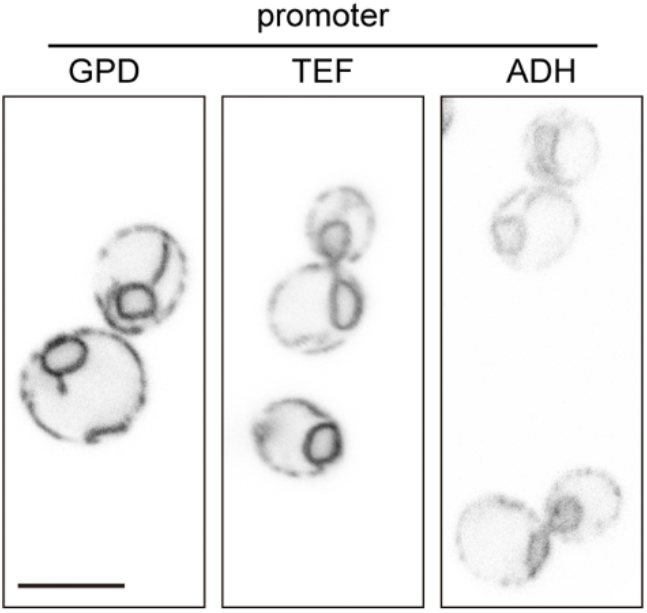
Localization of mNG-hRft1 expressed under control of promoters of different strength. The mNG-hRft1 construct was integrated into mCherry-HDEL-expressing wild-type cells, using three different promoters (GPD, TEF, ADH) to drive expression. The resulting cells (*P_GPD_*-mNG-hRft1 mCherry- HDEL, *P_TEF_*-mNG-hRft1 mCherry-HDEL, and *P_ADH_*-mNG-hRft1 mCherry-HDEL) were cultured in YPD medium to log-phase and imaged by confocal fluorescence microscopy to visualize mNG-hRft1. Images were taken under the same conditions, using the same microscope settings. Promoter strength GPD>TEF>ADH. Scale bar = 5 μm.

**Supplementary Figure S3.**
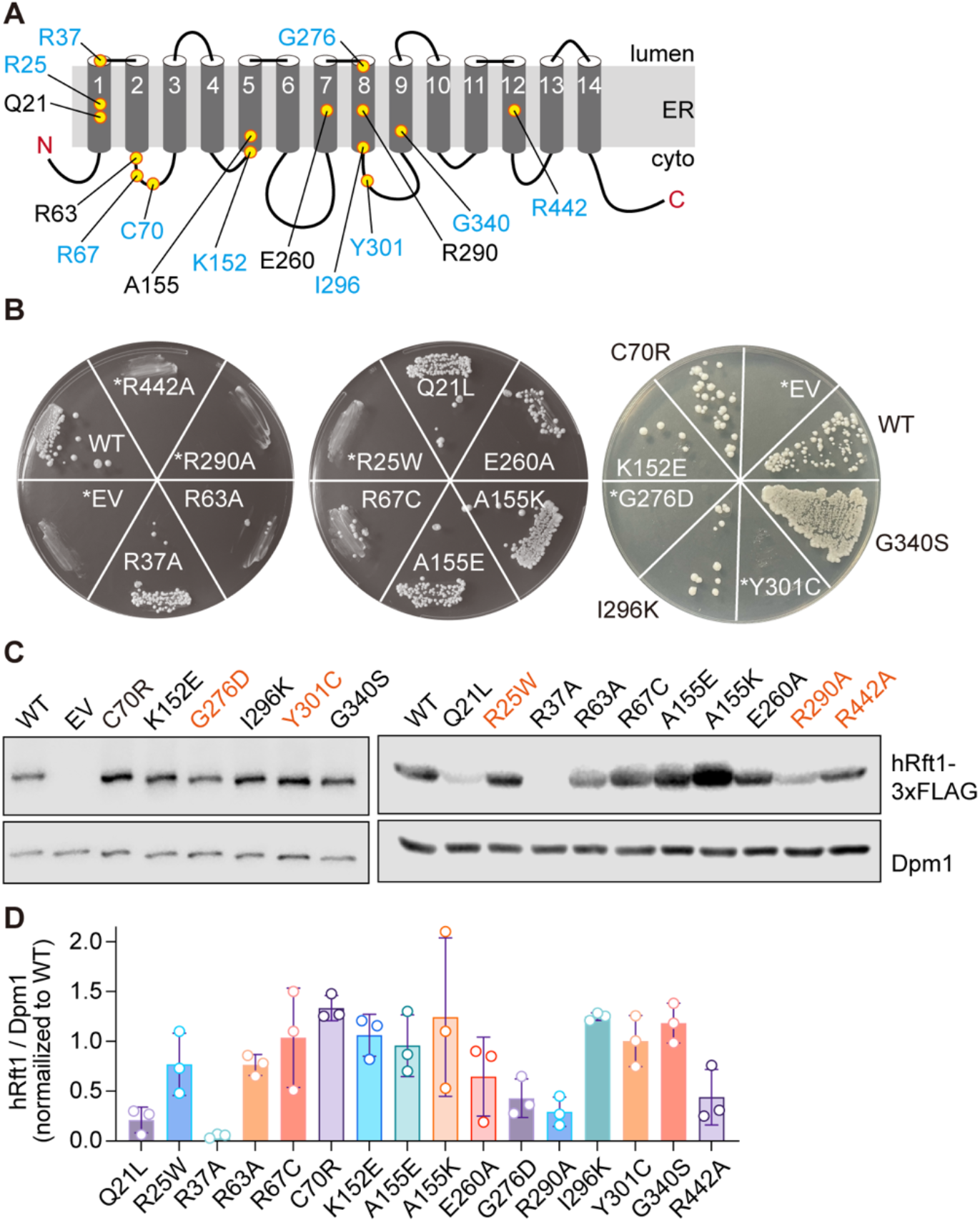
Restriction of hRft1 to the nuclear ER does not affect function. A. Topology model of hRft1 as shown in Figure 4A. Sites that were mutated in this study are indicated by yellow circles. Residues marked in blue correspond to Rft1-CDG mutations. B. KSY512 cells (rft1Δ←hRft1(URA3)) were transformed with HIS3 plasmids for expression of hRft1 point mutants with a C-terminal 3xFLAG tag. Colonies picked from SD(-Ura, -His) plates were streaked onto plates containing 5-FOA (1 mg/ml) to eject the *URA3* plasmid. Images were taken after 3 days of incubation at 30°C. The hRft1 mutants that did not grow on the 5-FOA plate are marked with an asterisk. EV, empty vector, WT, wild-type hRft1. C. Wild type (BY4741) cells carrying *HIS3* plasmids for expression of hRft1 point mutants with a C-terminal 3xFLAG tag were cultured in SD(-His) medium and harvested. Immunoblot assay was performed using anti-FLAG and anti-Dpm1 antibodies. D. hRft1-3xFLAG and Dpm1 (loading control) were quantified from panel C and two additional biological replicates. The ratio of the intensity of the hRft1 band to Dpm1 was calculated and normalized to that of the WT sample (mean ± S.D., n=3 biological replicates).

**Supplementary Figure S4.**
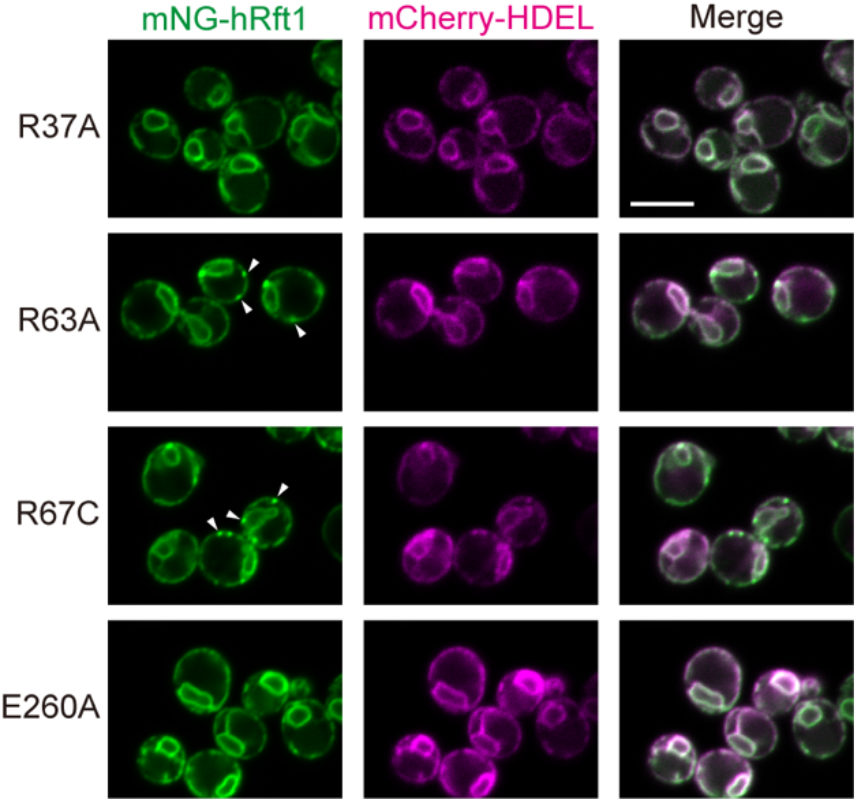
Subcellular localization of hRft1 point mutants. *P_GPD_*-mNG-hRft1^R37A/R63A/R67C/E260A^ mCherry-HDEL cells (YAKM248-251, Table 2) were cultured in YPD medium to log-phase and imaged by confocal fluorescence microscopy. The arrowheads indicate hRft1 varicosities seen in the cortical ER (2, 50, 54 and 25% of R37A, R63A, R67C and E260A-expressing cells, respecjvely (n>100 cells counted in each case) displayed these varicosijes). Scale bar = 5 μm.

